# Antibody-mediated prevention of vaginal HIV transmission is dictated by IgG subclass in humanized mice

**DOI:** 10.1101/2022.07.07.499107

**Authors:** Jacqueline M. Brady, Meredith Phelps, Scott W. MacDonald, Evan C. Lam, Adam Nitido, Dylan Parsons, Christine L. Boutros, Cailin E. Deal, Serah Tanno, Harini Natarajan, Margaret E. Ackerman, Vladimir D. Vrbanac, Alejandro B. Balazs

**Author notes:** Direct all correspondence to: Dr. Alejandro Balazs, Ragon Institute of MGH, MIT, and Harvard, 400 Technology Square, Cambridge, MA 02139.

## Abstract

HIV broadly neutralizing antibodies (bNAbs) are capable of both blocking viral entry and recruiting innate immunity to HIV-infected cells through their fragment crystallizable (Fc) region. Vaccination or productive infection results in a polyclonal mixture of class-switched IgG antibodies comprised of four subclasses, each encoding distinct Fc regions that differentially engage innate immune functions. Despite evidence that innate immunity contributes to protection, the relative contribution of individual IgG subclasses is unknown. Here we use vectored immunoprophylaxis (VIP) in humanized mice to interrogate the efficacy of individual IgG subclasses during prevention of vaginal HIV transmission by VRC07, a potent CD4-binding site directed bNAb. We find that VRC07-IgG2, which lacks Fc-mediated functionality, exhibits significantly reduced protection *in vivo* relative to other subclasses. However, even low concentrations of highly functional VRC07-IgG1 yields substantial protection against vaginal challenge, suggesting that interventions capable of eliciting modest titers of functional subclasses may provide meaningful benefit against infection.

## INTRODUCTION

Broadly neutralizing antibodies (bNAbs) target conserved regions of the HIV envelope glycoprotein (Env) to inhibit infection through both direct neutralization of virus as well as recruitment of innate immunity to HIV-infected cells. The fragment crystallizable (Fc) regions of antibodies engage Fc gamma receptors (FcɣRs) on the surface of innate immune cells to elicit antibody-dependent cellular cytotoxicity (ADCC), phagocytosis (ADCP), and other effector functions. The immunoglobulin gamma (IgG) isotype comprises the majority of class-switched antibody produced in response to either vaccination or pathogen infection (Janeway et al., 2001). However, IgG in humans is composed of four individual IgG subclasses (IgG1-4), each of which exhibit a unique pattern of binding to the numerous FcɣR proteins expressed by innate immune cells, yielding distinct capacities for each subclass to elicit effector functions (Vidarsson et al., 2014).

The antibody mediated prevention (AMP) study recently demonstrated that passive transfer of a CD4-binding site (CD4bs)-directed broadly neutralizing antibody (bNAb) of the IgG1 subclass prevented transmission of sensitive strains to participants (Corey et al., 2021). Given the remarkable diversity of circulating strains, significant effort is presently focused on the development of HIV vaccine regimens capable of eliciting bNAbs with maximal breadth and potency (Sok and Burton, 2018). However, existing vaccination regimens yield polyclonal antibody responses composed of all IgG subclasses (Visciano et al., 2012), which may result in sub-optimal protective efficacy. Although, patients who naturally control their HIV infections have been shown to harbor a greater proportion of functional IgG subclasses (Banerjee et al., 2010), the efficacy of each individual subclass during prevention of HIV transmission has not been defined. Studies in both non-human primates (NHPs) employing the SIV-HIV chimeric virus (SHIV) and humanized mice employing HIV have demonstrated that CD4 binding site (CD4bs)-directed bNAbs of the IgG1 subclass exhibit reduced protection when harboring Fc mutations that limit effector functions (Bournazos et al., 2014; Hessell et al., 2007). However, these findings have not held true for Fc-mutants of the V3-glycan-directed bNAb, PGT121, against either cell-associated or mucosal SHIV transmission in NHPs (Hangartner et al., 2021; Parsons et al., 2018). Despite these conflicting results, recent studies in macaques have shown that passive transfer of highly functional IgGs correlates with improved protection against SIV transmission (Alter et al., 2020). However, these studies were conducted in simian models which do not entirely recapitulate human FcɣR interactions (Crowley and Ackerman, 2019). In addition, while previous studies have examined the relationship between *in vitro* bNAb potency and *in vivo* dose necessary to prevent transmission of chimeric SHIV in the NHP model (Julg et al., 2017; Moldt et al., 2012a; Pegu et al., 2019), the precise relationship between bNAb concentration and risk of acquisition of transmitted-founder HIV in the context of a human immune system has not been established. The minimum protective dose of IgG bNAbs and effector cell activities that coordinate with distinct human IgG subclasses to prevent HIV transmission remain unknown, despite their critical importance to the success of ongoing efforts the develop vaccines and prophylactic interventions. A central issue has been that most animal models do not permit an examination of HIV infection in the context of a human immune response.

In this study, we utilize humanized mice, which harbor human innate immune cells bearing the full complement of FcɣRs, to determine the relative potential of each IgG subclass to prevent infection by a transmitted founder strain of HIV. For antibody delivery, we and others have described vectored immunoprophylaxis (VIP), using adeno-associated virus (AAV) vectors to deliver transgenes encoding antibodies to muscle tissue, resulting in long-term, systemic production of bNAbs capable of protecting humanized mice and NHPs from HIV or SIV challenge (Balazs et al., 2011, 2014). This dose-dependent expression of genetically encoded bNAb enables dissection of the precise contribution of each IgG subclass *in vivo*. Using this approach, our study defines the individual contributions of IgG subclasses during prevention of vaginal HIV transmission. Our findings suggest that subclasses with diminished capacity to mediate both ADCP and ADCC exhibit reduced protection as compared to more functional IgG subclasses, but only at low circulating antibody concentrations. This is particularly relevant to bNAb elicitation by vaccination where antibody levels are also expected be low. Analysis of the relative efficacy of protection across a range of IgG1 doses yields a statistical model of the relationship between bNAb concentration and risk of infection, demonstrating that even modest concentrations of functional antibodies can prevent transmission.

## RESULTS

### Generation and in vitro characterization of VRC07 IgG subclasses

To produce antibodies with different Fc functionality, we generated AAV serotype 8 (AAV8) expression vectors encoding the VRC07 variable region followed by the constant region of each human IgG subclass: IgG1, IgG2, IgG3, and IgG4. Heavy and light chains were expressed from a single transgene separated by a furin-2A picornavirus self-processing peptide as has been previously described (Balazs et al., 2011). As anticipated, Fc modulation of VRC07 had minimal impact on the ability of purified proteins to neutralize HIV *in vitro* (**Figure 1A**) or bind to HIV-1 Env expressed on the surface of cells (**Figure 1B**). However, Fc alteration did impact binding of FcɣRs to VRC07-bound target cells as measured by flow cytometry. FcɣRs bound to VRC07 IgG1 and IgG3 at lower half-maximal binding concentrations than the other VRC07 IgG subclasses, indicating a higher affinity binding interaction between these IgG subclasses and FcɣRs (**Figure 1C** and **S1**). Though IgG4 demonstrated comparable binding to FcɣR1a, it had markedly reduced binding to all other FcɣRs tested. Nearly all FcɣRs exhibited poor binding to IgG2, consistent with previous reports for other specificities (Brown et al., 2017).

**Figure 1.**
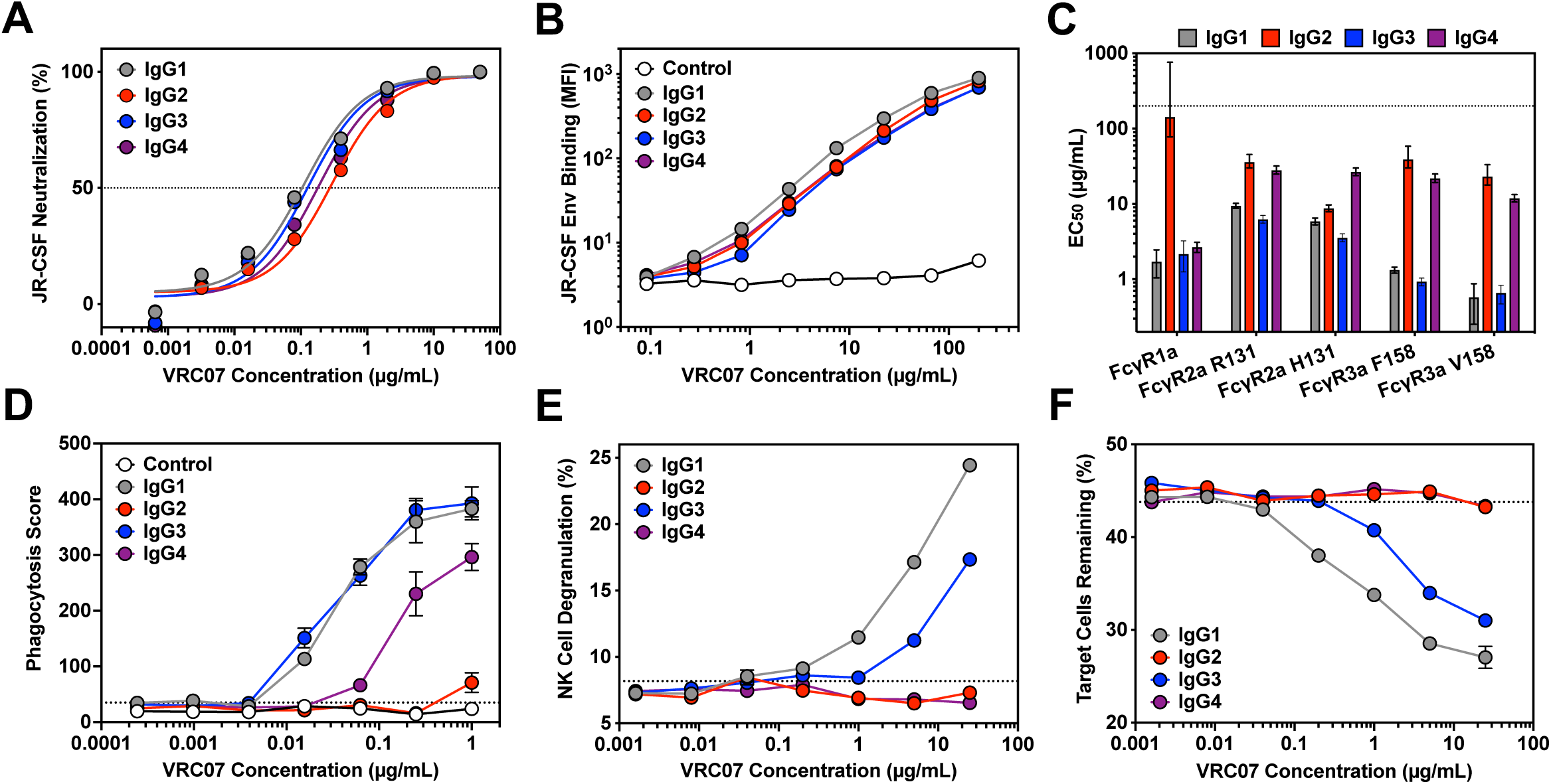
Impact of IgG subclass switching on the ability of VRC07 to bind FcɣRs and mediate effector function activity. **(A)** In vitro neutralization activity of purified VRC07 IgG subclasses against HIV_JR-CSF_ as measured by TZM-bl neutralization assays. Error bars represent SEM. **(B)** Binding of VRC07 IgG subclasses or a control antibody to HIV_JR-CSF_ Env-expressing target cells as measured by flow cytometry with a pan-IgG-Fc detection reagent. Error bars represent SEM. **(C)** Half-maximal binding concentration (EC50) of cell surface-bound VRC07 IgG subclasses to purified fluorescent-labeled FcɣR proteins as measured by flow cytometry. The dotted line represents the maximum concentration tested. Error bars denote the 95% confidence interval. **(D)** Phagocytosis of HIV_JR-CSF_ Env-expressing target cells by THP-1 cells mediated by VRC07 IgG subclasses or a non-HIV-specific control antibody as measured by flow cytometry. The dotted line indicates the average phagocytic score in the absence of antibody. Error bars represent SEM. **(E and F)** Ability of VRC07 IgG subclasses to mediate NK cell degranulation (E) or specific killing of HIV_JR-CSF_ Env-expressing target cells (F) as measured by flow cytometry. The dotted line indicates the average value in the absence of antibody. Error bars represent SEM.

To determine whether FcɣR binding differences impacted Fc function, we measured the ability of VRC07 IgG subclasses to mediate ADCP and ADCC activity in cell-based effector function assays. We began by measuring the ability of these antibodies to induce THP-1 cells to engulf CEM.NKr target cells stably expressing HIV_JR-CSF_ Env and a ZsGreen fluorescent reporter protein (**Figure 1D**). In this assay, IgG1 and IgG3 showed equivalent ability to facilitate phagocytosis by THP-1 effector cells. IgG4 led to reduced phagocytic activity compared to IgG1, but still maintained some activity at higher concentrations. IgG2, on the other hand, was unable to mediate any phagocytic activity over background. To confirm these findings, we also used an antigen-coated bead-based phagocytosis assay as previously described^22^. Interestingly, IgG3 mediated considerably more potent phagocytosis of HIV_JR-CSF_ gp120-coated beads than either IgG1 or IgG4 as has been previously observed for other HIV-specific monoclonal antibodies (Chu et al., 2020; Richardson et al., 2019). IgG2, however, mediated little to no phagocytic activity of beads over background even at the highest concentration tested (**Figure S2A**). To confirm the generalizability of these observations across HIV Env variants, we repeated these experiments using HIV_REJO.c_ gp120-coated beads as targets and observed similar trends (**Figure S2B**). However, when HIV_REJO.c_ Env-expressing target cells were used, IgG1, IgG3 and IgG4 exhibited comparable ability to mediate phagocytosis by THP-1 effector cells. IgG2 also facilitated detectable phagocytic activity at higher concentrations, though it was substantially lower than that observed for the other IgG subclasses (**Figure S2C**).

To measure ADCC, we performed analogous cell-based killing assays using the HIV_JR-CSF_ Env-expressing cells mixed with non-transduced, parental CEM.NKr cells. Incubation of the mixed target population with antibody and donor-derived NK cells led to a considerable increase in degranulation of NK cells which corresponded with specific depletion of Env-expressing target cells from the population. While this observation held true for IgG1 and IgG3 antibodies, neither IgG2 nor IgG4 antibodies were able to induce NK cell degranulation or target cell killing. Even at the highest concentration tested, these antibodies were indistinguishable from the control in which no antibody was added (**Figure 1E** and **1F**). When HIV_REJO.c_ Env-expressing target cells were used, we observed similar results (**Figure S2D** and **S2E**). Taken together, these assays demonstrate that changes to VRC07 IgG subclass have minimal impact on neutralization and epitope binding, but significantly alter the capacity of VRC07 to mediate Fc effector function activity, with IgG1 and IgG3 mediating both ADCC and ADCP, IgG4 mediating ADCP but no ADCC, and IgG2 mediating neither innate function.

### *In vivo* characterization of VRC07 IgG subclasses

We next determined the potential of the VRC07 IgG subclasses to mediate *in vivo* HIV protection when delivered via VIP. To this end, we performed an HIV challenge study in human peripheral blood mononuclear cell-engrafted NOD/SCID/ɣc^-/-^ (NSG) mice (huPBMC-NSG) using established conditions we previously described (Balazs et al., 2011) (**Figure 2A**). Importantly, the VRC07 IgG subclass variants exhibited no difference in neutralizing activity against the challenge strain, HIV_NL4-3_, *in vitro* (**Figure 2B**). To achieve *in vivo* expression of VRC07 IgG subclasses, we performed intramuscular injections of AAV8 encoding each antibody construct. Eight weeks after intramuscular (IM) AAV administration, mice achieved average plasma concentrations ranging from 50 to 80 µg/mL for VRC07 IgG1, IgG2 and IgG4. (**Figure 2C**). Mice expressing VRC07 IgG3 achieved plasma concentrations of approximately 10 µg/mL, which is significantly above the 5 µg/mL previously shown to be the minimum protective dose for VRC07 IgG1 against HIV_NL4-3_ in this model (Balazs et al., 2014). Following 8 weeks of expression, the NSG mice were humanized via injection of activated human PBMCs and two weeks later were challenged with a single intravenous (IV) administration of HIV_NL4-3_ (**Figure 2A**). HIV infection was assessed by weekly quantitation of both CD4^+^ T cell depletion and viral load in peripheral blood (**Figure 2D**). Seven of the 9 control mice that received AAV8-luciferase became infected following challenge whereas all mice that received AAV8-VRC07, regardless of the IgG subclass, were completely protected (**Figure 2E**). These results demonstrate that all four IgG subclasses are capable of mediating protection *in vivo* when present at high plasma concentration in a simplified humanized mouse model.

**Figure 2.**
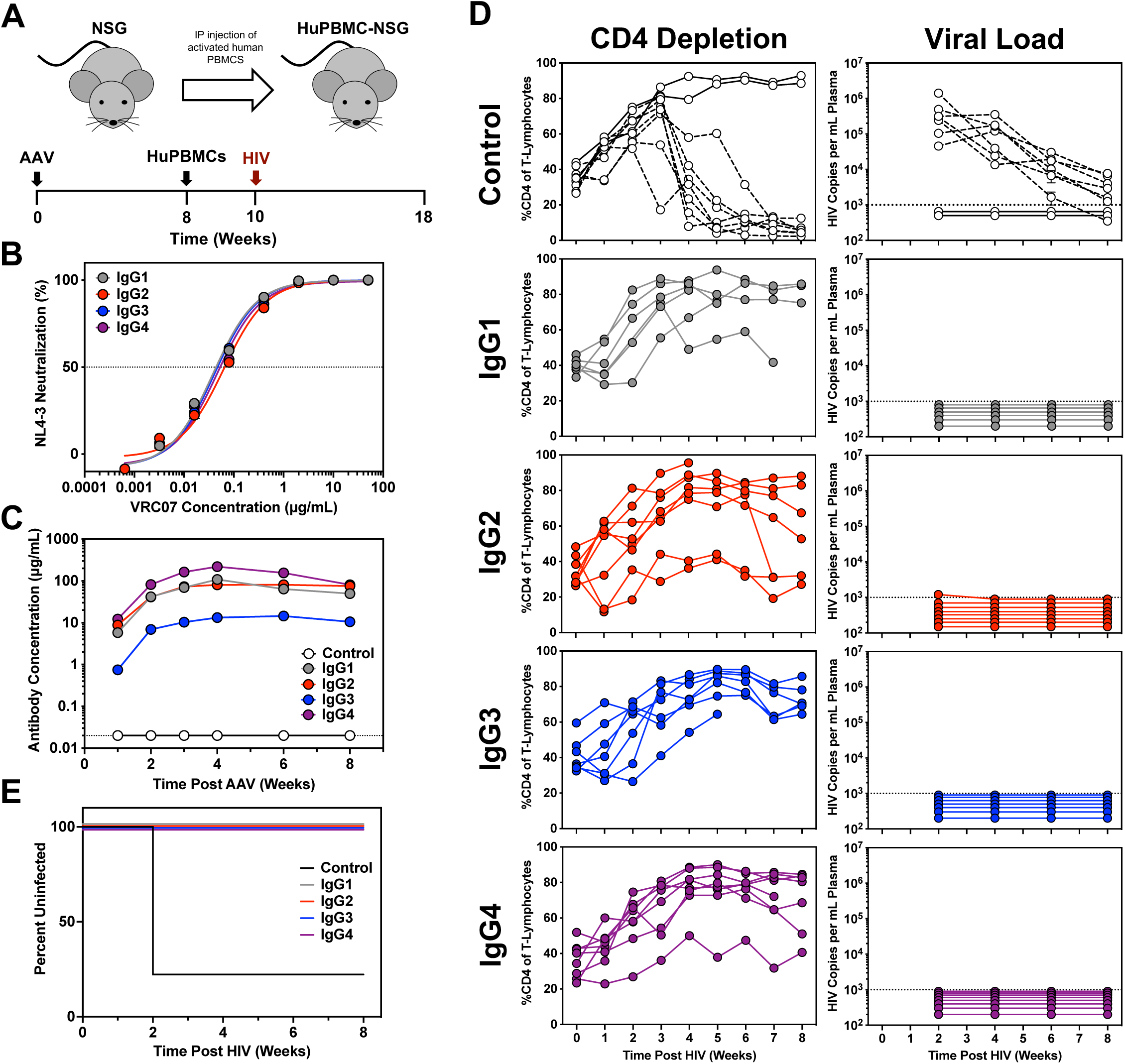
Impact of IgG subclass switching on the ability of VRC07 to protect from HIV challenge in huPBMC-NSG mice. **(A)** Overview of the huPBMC-NSG mouse model with a schematic representation of the experimental timeline. **(B)** In vitro neutralization activity of purified antibodies against the challenge strain, HIV_NL4-3_, as measured by a TZM-bl neutralization assay. Error bars represent SEM. **(C)** Average plasma antibody concentrations achieved in NSG mice following IM injection of 2.5 x 1011 GC of AAV expressing a given VRC07 IgG subclass or luciferase as a negative control (n=7-11 mice per group). Error bars represent SEM. **(D)** CD4 depletion (left) and viral load (right) in the peripheral blood of huPBMC-NSG mice expressing luciferase or a VRC07 IgG subclass following IV challenge of 280 TCID50 HIV_NL4-3_. Viral load was measured by qPCR. The dotted lines indicate the limit of detection, 1000 copies per mL. CD4 depletion was measured by flow cytometry. The dashed lines indicate mice that were confirmed to be infected by qPCR. **(E)** Kaplan-Meier survival curves for huPBMC-NSG mice expressing the given VRC07 IgG subclass or negative control.

### Efficacy of VRC07 IgG subclasses against vaginal HIV transmission

As most HIV infections occur through sexual transmission, we sought to investigate the impact of IgG subclass on VRC07-mediated protection in this context using the BLT humanized mouse model. This model consists of surgically implanted human thymic tissue and transplantation of genetically matched CD34^+^ stem cells, resulting in extensive multi-lineage engraftment of human immune cells throughout the animal, which render it susceptible to vaginal or rectal challenge (Balazs et al., 2014; Denton et al., 2008; Stoddart et al., 2011; Sun et al., 2007) (**Figure 3A**).

**Figure 3.**
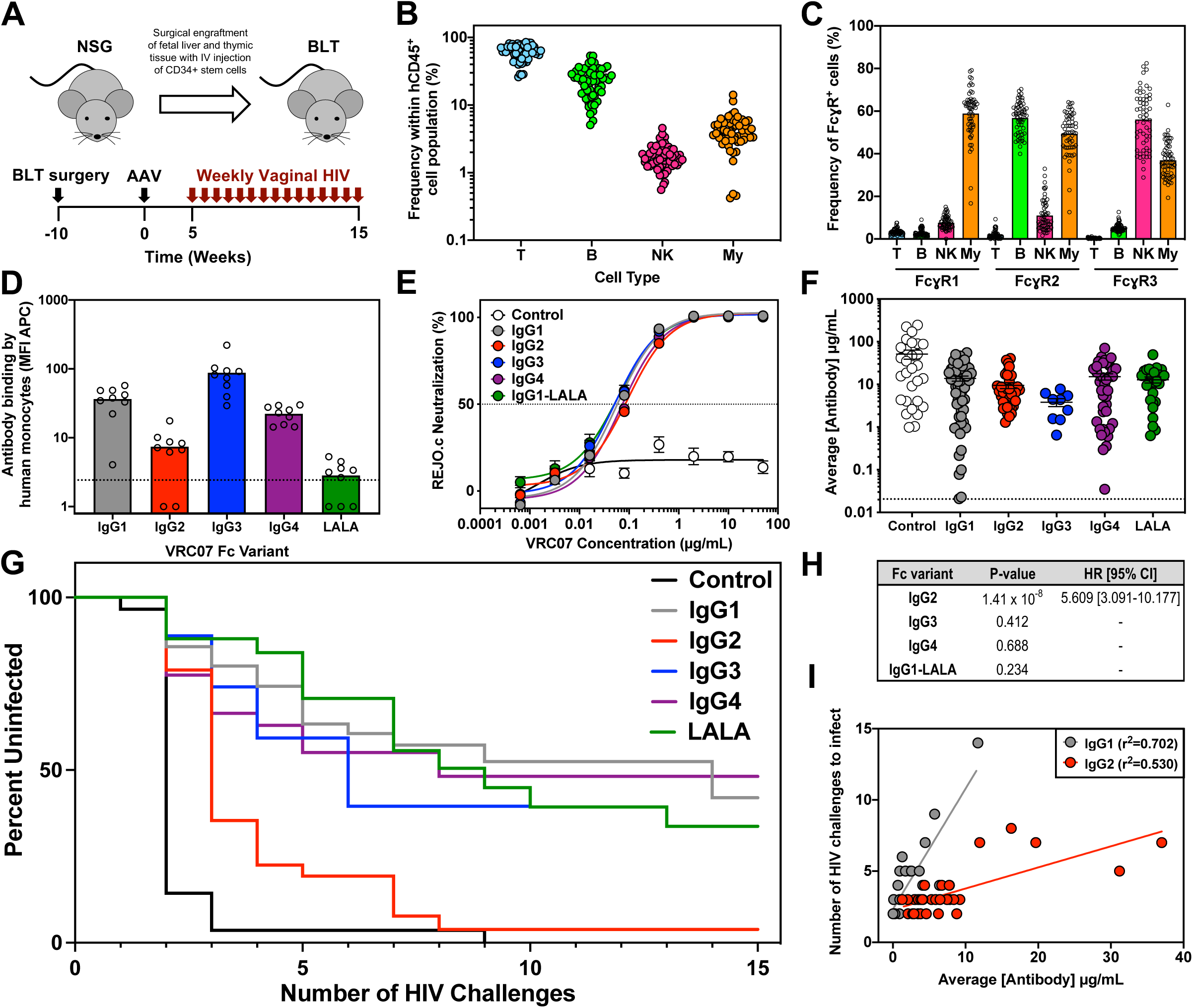
Protective efficacy of VRC07 IgG subclasses against repetitive HIV vaginal challenge in BLT humanized mice. **(A)** Overview of BLT humanized mouse model and schematic representation of experimental timeline. **(B)** The frequency of human T cells (CD45+, CD3+), B cells (CD45+, CD19+), NK cells (CD45+, CD3-, CD19-, HLA-DR-, CD56+), and myeloid cells (CD45+, CD3-, CD19-, CD11c+, HLA-DR+) in the peripheral blood of uninfected BLT mice (n=60) 10 weeks post surgical engraftment as measured by flow cytometry. **(C)** Percentage of the human immune cell subsets from panel (B) expressing human FcɣR1, FcɣR2, or FcɣR3 as measured by flow cytometry. Bars represent the mean. **(D)** Binding of AF647-labeled VRC07 Fc variants by human monocytes (CD45+, CD3-, CD56-, CD14+) isolated from the spleens of BLT mice (n=9) across two separate BLT batches as measured by flow cytometry. The dotted line indicates the average MFI in the absence of antibody. Bars represent the mean. **(E)** In vitro neutralization activity of purified VRC07 IgG subclasses against the challenge strain, HIV_REJO.c_, as measured by a TZM-bl neutralization assay. **(F)** Average plasma antibody concentrations achieved in BLT mice from the start of vaginal challenges through HIV acquisition or death. Mice were given 1.0 x 10^10^ – 2.5 x 10^11^ GC of AAV expressing the given antibody and plasma concentrations were measured weekly by ELISA. Graphs represent combined data from three separate BLT experiments with the exception of VRC07-IgG3 (1 experiment) and VRC07-LALA (2 experiments). **(G)** Kaplan-Meier survival curves for mice expressing different VRC07 IgG subclasses. **(H)** Cox regression analysis was used to evaluate the effect of Fc modulation on the rate of HIV acquisition across multiple BLT experiments while controlling for antibody concentration. A hazard ratio (HR) and 95% confidence interval (CI) are only reported if the p-value was less than 0.05. **(I)** Relationship between the average circulating antibody concentration (µg/mL) and the number of HIV challenges required for infection to occur over the course of challenge in BLT humanized mice. Lines represent the result of a linear regression for mice expressing VRC07 IgG1 or VRC07 IgG2.

To determine the potential for human cells in BLT mice to participate in innate immune interactions with human anti-HIV antibodies, we first characterized the relative frequency of innate immune cell types in peripheral blood. We noted that BLT mice harbored T cells, B cells, and myeloid cells at levels similar to those found in human blood (Yi et al., 2019), while NK cells were present at lower frequencies (**Figure 3B**). Flow cytometric analysis of immune cells in the peripheral blood of uninfected BLT humanized mice revealed that NK and myeloid cell subsets expressed the expected pattern of human FcɣRs (**Figure 3C**). Specifically, a large proportion of circulating NK cells expressed FcɣR3 (CD16), the receptor involved in mediating ADCC. Furthermore, a large proportion of the myeloid cell subset, which included monocytes and dendritic cells, also expressed FcɣR3 in addition to FcɣR2 (CD32) and FcɣR1 (CD64) which are involved in mediating ADCP.

To determine the potential of FcɣR-expressing cells in BLT mice to interact with the four subclasses of VRC07, we performed additional flow cytometry using fluorescently labeled VRC07 IgG1-IgG4 proteins. As a negative control, we included a VRC07 IgG1 variant with two leucine to alanine substitutions at positions 234 and 235 (VRC07 LALA), which has been previously shown to exhibit reduced binding to human FcɣRs (Hezareh et al., 2001). Exposure of splenocytes harvested from BLT mice to fluorescently labeled VRC07 proteins revealed that monocytes interacted most strongly with IgG1 and IgG3. IgG4 exhibited modest staining whereas IgG2 and LALA resulted in limited interaction with monocytes (**Figure 3D** and **S3**). Interestingly, NK cells in the spleen did not interact appreciably with labeled VRC07 in this experiment, however we observed a similar lack of NK cell staining in one of two human donors, suggesting that human immune cells in BLT mice display similar patterns of interactions with IgG subclasses as immune cells in humans (**Figure S3**).

Importantly, IgG subclass did not impact the *in vitro* neutralizing activity of these antibodies against the transmitted founder challenge stain, HIV_REJO.c_ (**Figure 3E**). We included VRC07 IgG1 harboring the LALA mutation in these studies as this Fc variant has previously been used to assess the contribution of effector function to bNAb-mediated protection. In effector function assays analogous to those described above, VRC07 LALA exhibited reduced binding to all FcɣRs compared to VRC07 IgG1 and failed to mediate NK cell degranulation or killing of Env-expressing target cells. However, VRC07 LALA retained some ability to mediate phagocytosis by THP-1 effector cells (**Figure S4**).

To determine the relative protective activity of each IgG subclass against repetitive challenge with HIV, we administered AAV8 expressing each VRC07 subclass (IgG1, IgG2, IgG3, IgG4, or LALA) to BLT mice at different doses to achieve plasma antibody concentrations ranging from 70 µg/mL to less than 0.05 µg/mL (**Figure 3F** and **S5**). Five weeks after AAV administration, we initiated weekly vaginal challenges of HIV_REJO.c_ for 15 consecutive weeks, and measured the rate of infection by plasma viral load (**Figure S6**).

Interestingly, VRC07 IgG3, IgG4, and LALA did not exhibit significantly different *in vivo* efficacy as compared to IgG1 (**Figure 3G**). Overall, VRC07 IgG3 achieved lower plasma antibody concentrations than the other Fc variants, with mice expressing less than 1.5 µg/mL exhibiting no protection and those expressing over 4 µg/mL exhibiting either delayed acquisition, or complete protection. Mice expressing VRC07 IgG4 exhibited a wide range of antibody concentrations from less than 1 µg/mL to over 70 µg/mL. Most animals expressing less than 1 µg/mL VRC07 IgG4 exhibited no protection while those expressing above 5 µg/mL of VRC07 IgG4 exhibiting delayed acquisition or complete resistance. Interestingly, mice expressing VRC07 LALA followed the same trend with those harboring less than 2 µg/mL in circulation exhibiting no protection while those above 6 µg/mL exhibiting a substantial delay in HIV acquisition or complete protection. Of note, all of these IgG subclasses including IgG1-LALA retained substantial phagocytic activity *in vitro*, particularly against cells expressing REJO.c envelope matching the challenge strain used in this experiment (**Figure S2 and S4**).

In contrast, VRC07 IgG2 showed a significant reduction in protective efficacy when compared to IgG1. Despite average plasma IgG2 concentrations between 1.2 and 9.2 µg/mL, 28 mice became infected within 4 challenges, while nearly all others were infected within 8 challenges. The only mouse to achieve complete protection from 15 repeated vaginal challenges expressed an average circulating VRC07 IgG2 concentration over 40 µg/mL. When all Fc variants were compared to IgG1 using a Cox regression analysis, we found a statistically significant association (p=1.41x10^-8^) between IgG2 and the rate of HIV acquisition (**Figure 3H**).

The associated hazard ratio indicates that mice expressing VRC07 IgG2 had greater than 5.6-fold increased risk of HIV infection compared to mice expressing VRC07 IgG1, irrespective of antibody concentration. When directly comparing VRC07 IgG2-expressing mice to VRC07 IgG1-expressing mice within the same range of concentrations, the correlation between antibody concentration and the number of challenges required to infect were distinct (**Figure 3I**). These results demonstrate that significantly more IgG2 antibody is required to achieve the same reduction in risk provided by IgG1.

### Minimum protective dose of VRC07 IgG1 against vaginal HIV challenge in humanized mice

Given the extensive clinical use of IgG1 subclass antibodies in human passive transfer studies, we sought to better define the protective efficacy of this subclass. Our laboratory has previously shown that the G54W variant of VRC07 IgG1 can fully protect BLT mice from repeated challenges of the transmitted-founder strain, HIV_REJO.c_, when present at approximately 100 µg/mL in plasma (Balazs et al., 2014). To determine the minimum protective concentration of VRC07-IgG1, we analyzed HIV acquisition in BLT mice expressing a wide range of concentrations (**Figure 4A**). In addition, as a negative control, we compared these to BLT mice expressing 2A10, an irrelevant malaria-specific human IgG1 antibody (Deal et al., 2014). Five weeks after AAV administration, BLT mice received weekly vaginal challenges of HIV_REJO.c_ for 15 consecutive weeks, and infection was detected by plasma viral load (**Figure 4B**). To ensure reproducibility of the observed results and control for variable immune reconstitution, we combined results from three separate experiments performed with distinct batches of BLT mice.

**Figure 4.**
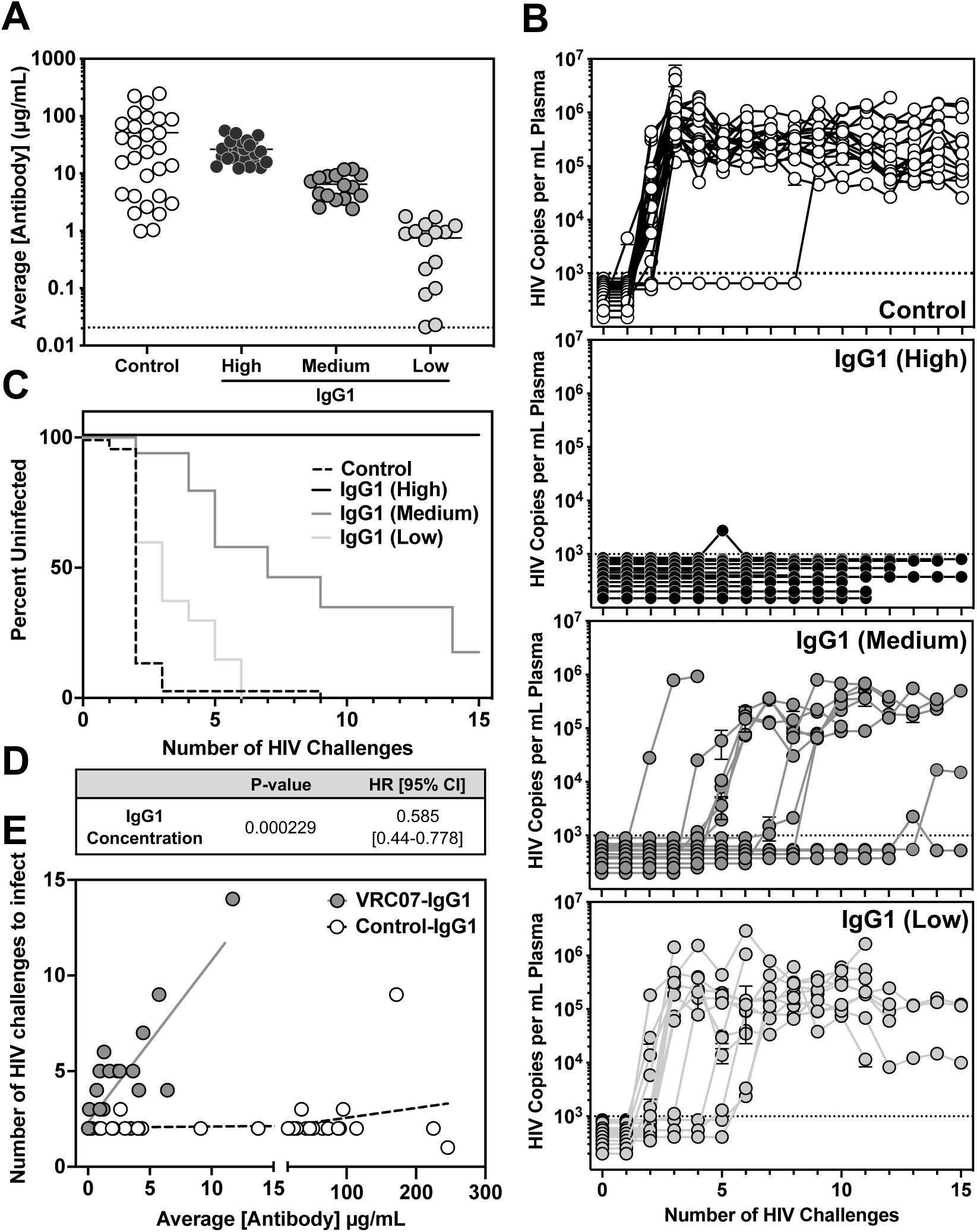
Protective efficacy of VRC07 IgG1 against repetitive HIV vaginal challenge in BLT humanized mice. (A) Average plasma antibody concentrations achieved in BLT mice from the start of vaginal challenge through HIV acquisition or death. Mice were grouped based on the average plasma VRC07 concentration achieved over the course of challenge. High – Greater than 12 µg/mL, Medium – 2-10 µg/mL, Low – Less than 2 µg/mL. The graphs represent combined data from three separate BLT experiments. (B) Viral load in the peripheral blood of BLT mice as measured by qPCR. The dotted line indicates the limit of detection (1000 copies/mL) for the assay. (C) Kaplan-Meier survival curves for mice expressing a control IgG1 antibody or varying concentrations of VRC07 IgG1. (D) Cox regression analysis was used to evaluate the effect of VRC07 IgG1 concentration on the rate of HIV acquisition across multiple BLT experiments. HR - hazard ratio, CI - confidence interval. (E) Relationship between the average circulating antibody concentration (µg/mL) and the number of HIV challenges required for HIV infection to occur over the course of challenge in BLT humanized mice. Lines represent the result of a linear regression for mice expressing VRC07 or 2A10 IgG1.

Across 29 mice expressing the 2A10 negative control antibody, we detected plasma antibody concentrations ranging from roughly 1 to 250 µg/mL. All but one of these mice, regardless of circulating antibody levels, became infected after 3 repeated vaginal challenges with HIV. The remaining control mouse became infected after 9 challenges (**Figure 4B** and **4C**). Mice receiving AAV8-VRC07-IgG1 were grouped based on average weekly plasma antibody concentration over the course of HIV challenge (**Figure S5**). This metric was used to categorize mice into high (greater than 12 µg/mL), medium (2-12 µg/mL) or low (less than 2 µg/mL) expression groups. All 25 mice in the high expression group were completely protected from 15 consecutive vaginal challenges with HIV_REJO.c_ (**Figure 4B** and **4C**). Within the medium group, nearly all mice expressing at least 6.5 µg/mL of VRC07 were fully protected, with a single exception that became infected after 14 repeated challenges. Remarkably, even mice with circulating antibody concentrations as low as 3-5 µg/mL exhibited delays in HIV acquisition, becoming infected only after 5 or more repeated challenges. In contrast, mice expressing less than 1 µg/mL typically became infected after 3 repeated challenges, similar to control mice expressing 2A10. However, two mice expressing between 0.95 and 1.78 µg/mL delayed HIV acquisition until 5 or 6 repeated challenges.

Overall, these findings are consistent with a dose-dependent effect of VRC07-IgG1 against HIV_REJO.c_ challenge, with even relatively low plasma concentrations affording robust protection from vaginal transmission of this sensitive strain. A Cox regression analysis performed across this range of antibody concentrations found a statistically significant association (p=0.000229) between the rate of HIV acquisition and VRC07-IgG1 concentration with a hazard ratio of 0.585, indicating that every 1 µg/mL increase in circulating antibody concentration results in a 1.71-fold decreased risk of HIV infection (**Figure 4D**). In contrast, animals expressing 2A10-IgG1 as a negative control exhibited no relationship between antibody concentration and the number of challenges required to become infected (**Figure 4E**).

## DISCUSSION

By switching IgG subclasses and site-directed mutagenesis we generated VRC07 IgG variants with differing capacities to elicit Fc-dependent functions. Despite their similarity in binding cell-surface Env and neutralizing viral particles, these changes significantly altered the affinity of VRC07 for Fc gamma receptors. As a consequence, VRC07 subclass variants demonstrated markedly different ability to mediate ADCC and ADCP functions *in vitro* as measured in cell-based assays.

In an effort to deconvolute the contribution of individual IgG subclasses during prophylaxis, we assessed the ability of each subclass to mediate protection *in vivo*. While all subclasses demonstrated the capacity for preventing HIV infection when present at high concentration during intravenous challenge, we found that VRC07 IgG2 exhibited significantly reduced protection against vaginal challenge in BLT humanized mice. While some BLT mice expressing VRC07-IgG2 exhibited delays in HIV acquisition, substantially higher plasma antibody concentrations were necessary to achieve this outcome as compared with other subclasses. Interestingly, VRC07-IgG2 was the only subclass unable to mediate either ADCP or ADCC activity *in vitro*. Our findings support a model in which neutralization is sufficient to mediate protection at high circulating antibody concentrations, while Fc-dependent functions contribute to antibody efficacy near the minimum protective dose. This holds particular relevance for vaccine efforts where, if successful, the elicitation of bNAbs will likely occur at low titer.

In our studies both IgG4 and IgG1-LALA conferred *in vivo* protection comparable to that of IgG1, suggesting that ADCC activity is insufficient to explain the differences observed in our model. Other studies in macaques have shown no appreciable differences in bNAb efficacy following NK cell-depletion (Parsons et al., 2018). Furthermore, a b12 variant with enhanced FcɣR3a binding and ADCC activity failed to improve protection against mucosal SHIVSF162P3 challenge in rhesus macaques (Moldt et al., 2012b). Notably, VRC07-IgG2 was the only subclass entirely lacking phagocytic activity *in vitro*. IgG2, IgG4, and LALA all exhibited a substantial reduction in affinity for FcɣR2a compared to VRC07 IgG1. However, IgG2 was the only Fc variant also lacking FcɣR1a binding. As a result, IgG4 and LALA maintained some ability to mediate phagocytosis of Env-expressing target cells. Furthermore, both IgG4 and LALA conferred protection against vaginal HIV challenge similar to more functional subclasses, suggesting that ADCP may be crucially important during antibody-mediated prevention of HIV transmission.

In support of this hypothesis, human clinical trials and NHP studies have also suggested ADCP as a potential mechanism of HIV prevention with other vaccine modalities (Shangguan et al., 2021). The HVTN 505 vaccine trial lacked overall efficacy, but found that ADCP by monocytes correlated with decreased HIV risk (Neidich et al., 2019). Similarly, IgG-driven monocyte-mediated phagocytosis correlated with reduced risk of SIV infection (Ackerman et al., 2018). Furthermore, an adjuvant enhancing antibody-dependent monocyte and neutrophil-mediated phagocytic responses augmented protection against stringent mucosal SHIV challenge in rhesus macaques (Om et al., 2020).

Although we do not know the immune cell subset(s) responsible for driving ADCP activity in BLT humanized mice, we have demonstrated that human monocytes in this model express Fc gamma receptors and differentially bind to VRC07 IgG subclasses. A study of human biopsy samples found FcɣR-expressing macrophages were highly enriched in most tissues and FcɣR-expressing neutrophils mediated the most efficient phagocytosis (Sips et al., 2016). This study also found negligible FcɣR-expressing NK cells in the female reproductive mucosa and other vulnerable tissues, suggesting that phagocytosis may be more likely to contribute to prevention (Sips et al., 2016). Additional studies will be necessary to dissect the precise effector mechanisms and immune cell populations contributing most to bNAb-mediated protection.

Our laboratory has previously shown that a variant of VRC07-IgG1 protected BLT mice from 21 repeated HIV_REJO.c_ challenges when expressed at nearly 100 µg/mL in circulation (Balazs et al., 2014). In the present study, we found that wildtype VRC07-IgG1 concentrations 8-fold lower than previously published still achieved complete protection from 15 consecutive HIV_REJO.c_ challenges. As little as 5 µg/mL of VRC07-IgG1 in circulation substantially delayed HIV acquisition in BLT mice, suggesting that small amounts of functional bNAbs could have meaningful impact on the transmission of neutralization-sensitive HIV strains. This is particularly relevant in light of progress made in the clinical translation of VIP to humans, given reports of durable expression of µg/mL levels of VRC07-IgG1 in patients after vector administration in the VRC603 study (Casazza et al., 2021).

Despite a half-maximal inhibitory concentration (IC_50_) of VRC07 against HIV_REJO.c_ of 0.05 µg/mL *in vitro*, our *in vivo* challenge studies demonstrated that all mice expressing greater than 12 µg/mL of VRC07 were completely protected from vaginal HIV_REJO.c_ challenge. This observation suggests that optimal efficacy *in vivo* requires a VRC07-IgG1 plasma concentration approximately 240 times higher than the *in vitro* IC_50_. However, our study employed a stringent challenge that led to infection of nearly all control mice after 1-2 challenges. In contrast, human heterosexual transmission is estimated to occur at a frequency between 1 in 100 and 1 in 1000 exposures (Hughes et al., 2012; Shattock and Moore, 2003). Therefore, our challenge regimen may reflect a high bar for antibody-mediated protection, possibly overestimating the concentration necessary for complete efficacy.

Overall, our findings support the conclusion that Fc subclass contributes significantly to bNAb-mediated HIV prevention through its role modulating antibody-dependent innate effector functions. This observation is in agreement with mucosal challenge studies in rhesus macaques in which LALA exhibited reduced b12 antibody activity against vaginal SHIVSF162P3 challenge (Hessell et al., 2007). However, more recent work reported that Fc-dependent functions did not contribute to the protective efficacy of PGT121 in pigtail macaques, where PGT121-LALA completely protected animals from intravenous challenge with splenocytes from a SHIVSF162P3-infected donor animal (Parsons et al., 2018). Even at lower circulating antibody concentrations, PGT121 and PGT121-LALA demonstrated comparable protective efficacy against high-dose vaginal challenge of SHIVSF162P3 (Hangartner et al., 2021). Given the results of our study, it will be essential that future experiments investigating the role of Fc function in bNAb efficacy compare Fc variant antibodies at a range of circulating antibody concentrations encompassing the minimum protective dose for the specific model being employed.

In conclusion, our findings suggest that IgG subclass is important for optimal prophylactic intervention against HIV, particularly at low bNAb concentrations. A maximally effective HIV antibody will likely be capable of broad, potent neutralization as well as significant effector function activity to maximize its potential to prevent infection at lower concentrations. Future work defining the precise effector mechanisms involved in bNAb-mediated protection will be crucial to the design of optimally effective antibody-based interventions.

## LIMITATIONS OF THE STUDY

The main focus of this study was to determine the role of IgG subclass during prevention of HIV transmission. For this purpose, we utilized several humanized mouse models which may not entirely recapitulate the immune environment of mucosal tissues in humans, possibly altering the efficiency of HIV transmission in the model and impacting the estimate of minimum protective dose for a given antibody. In particular, the relative scarcity of NK cells in the BLT model as compared to humans may underestimate the role of ADCC during antibody-mediated prevention. In addition, vectored delivery and production from cells other than B cells may result in antibodies with different glycosylation patterns than those elicited naturally, which may influence their function. Finally, the use of a challenge regimen resulting in productive HIV infection at a rate significantly above that seen in humans may underestimate the capacity for bNAbs to prevent infection, resulting in an overestimation of the minimum protective dose.

## ACKNOWLEDGEMENTS, FUNDING SUPPORT

We wish to thank Daniel Lingwood, PhD and Kiera Clayton, PhD for many useful and insightful discussions. H.N and M.E.A are supported by NIGMS and NIAID R01AI131975. J.M.B. is supported by Ruth L. Kirschstein Predoctoral Individual National Research Service Award 1F31AI131747-01A1, A.B.B. is supported by the National Institutes for Drug Abuse (NIDA) Avenir New Innovator Award DP2DA040254, the MGH Transformative Scholars Program as well as funding from the Charles H. Hood Foundation. This independent research was supported by the Gilead Sciences Research Scholars Program in HIV.

## AUTHOR CONTRIBUTIONS

J.M.B., M.P. and A.B.B. designed the experiments. J.M.B, M.P., S.W.M., E.C.L., A.N., D.P., C.L.B., and C.E.D carried out experiments and analyzed data. H.N. and M.E.A. offered suggestions for experiments and provided materials. S.T. and V.D.V. provided humanized mice. J.M.B. and A.B.B. wrote the paper with contributions from all authors.

## DECLARATIONS OF INTEREST

A.B.B. is a named inventor on patent US9527904B2 held by the California Institute of Technology describing the vector used in this study. The other authors declare that they have no competing interests.

## SUPPLEMENTAL FIGURE TITLES AND LEGENDS

**Figure S1.**
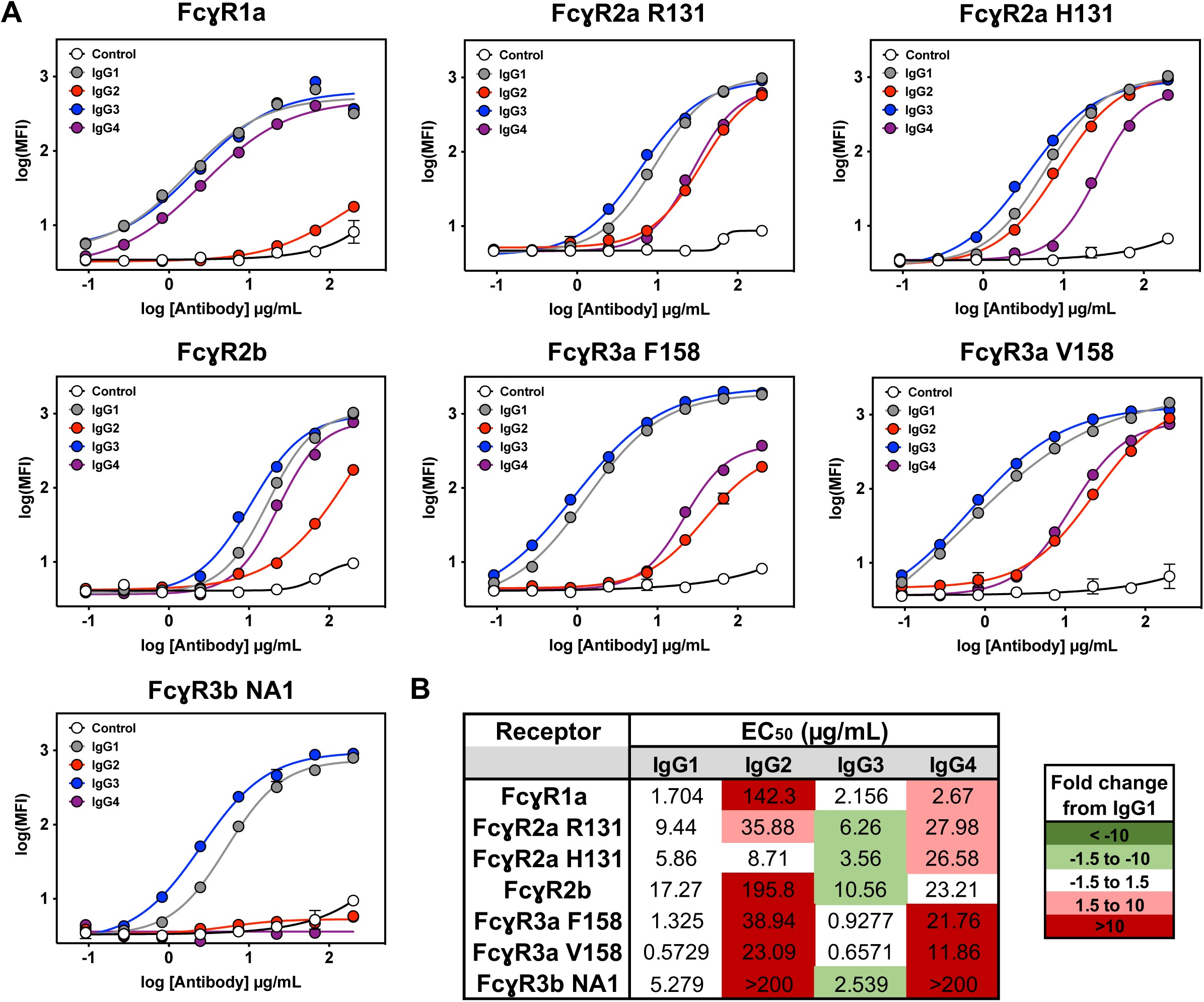
Impact of IgG subclass switching on the ability of VRC07 to bind purified FcɣRs, related to Figure 1. **(A)** FcɣR binding profiles for VRC07 IgG subclasses bound to HIV_JR-CSF_ Env-expressing CEM.NKr cells as measured by flow cytometry. The control is a non-HIV-specific IgG1 antibody. Error bars represent SEM. **(B)** EC_50_, or half-maximal binding concentrations (µg/mL) calculated for each VRC07 IgG subclass and FcɣR combination. The color of the box indicates the fold change in half-maximal binding compared to VRC07 IgG1.

**Figure S2.**
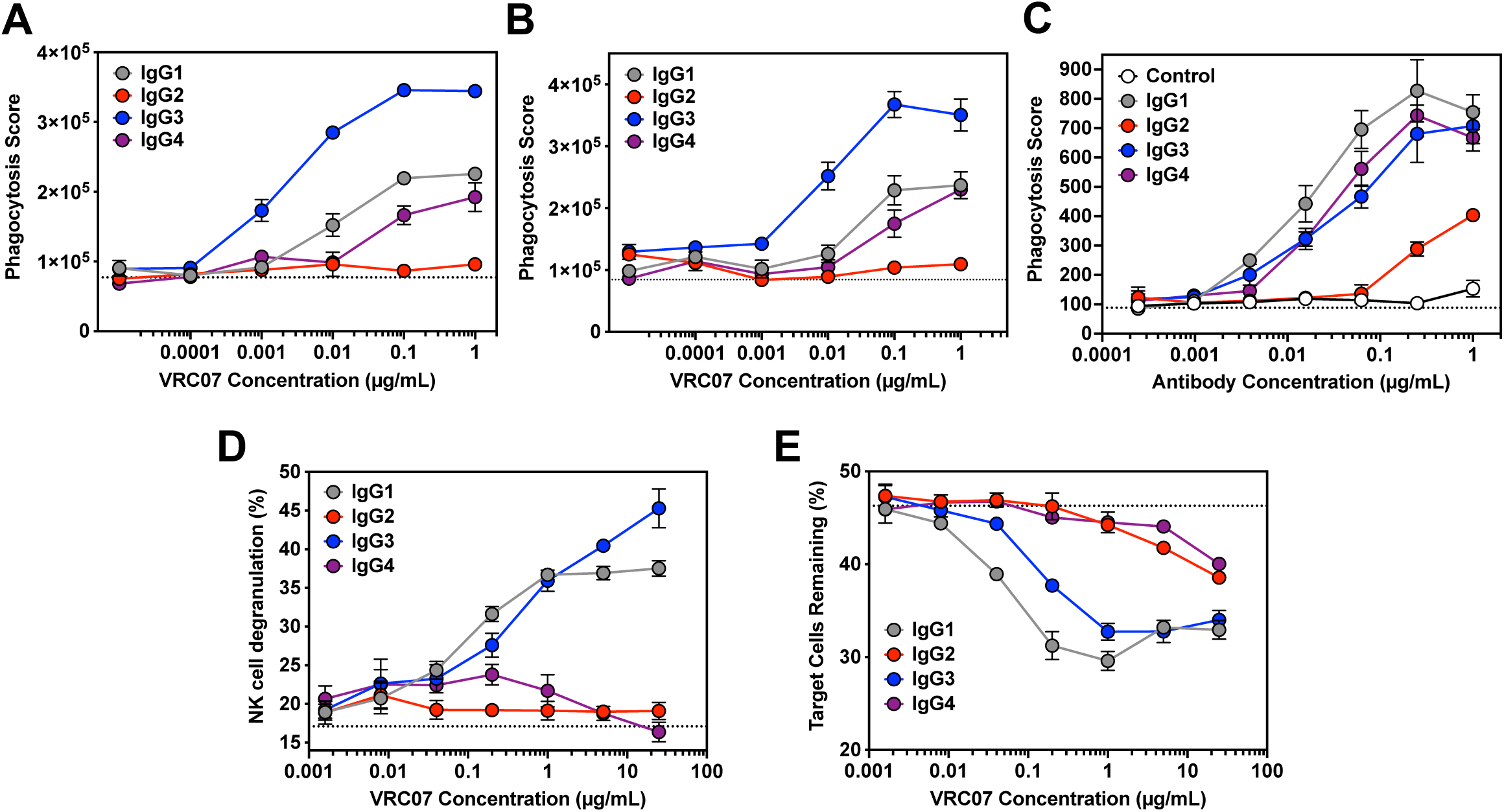
Impact of Fc subclass on the ability of VRC07 to mediate phagocytosis by THP-1 cells, related to Figure 1. **(A, B and C)** Phagocytosis of HIV_JR-CSF_ **(A)** or HIV_REJO.c_ **(B)** gp120-coated fluorescent beads, or HIV_REJO.c_ Env-expressing CEM.NKr target cells **(C)** by THP-1 cells in the presence of different VRC07 IgG subclasses as measured by flow cytometry. The dotted line indicates the average phagocytic score in the absence of antibody. **(D and E)** Ability of VRC07 IgG subclasses to mediate NK cell degranulation **(D)** as well as specific killing of HIV_REJO.c_ Env-expressing target cells **(E)** as measured by flow cytometry. The dotted line indicates the average value in the absence of antibody. Error bars represent SEM.

**Figure S3.**
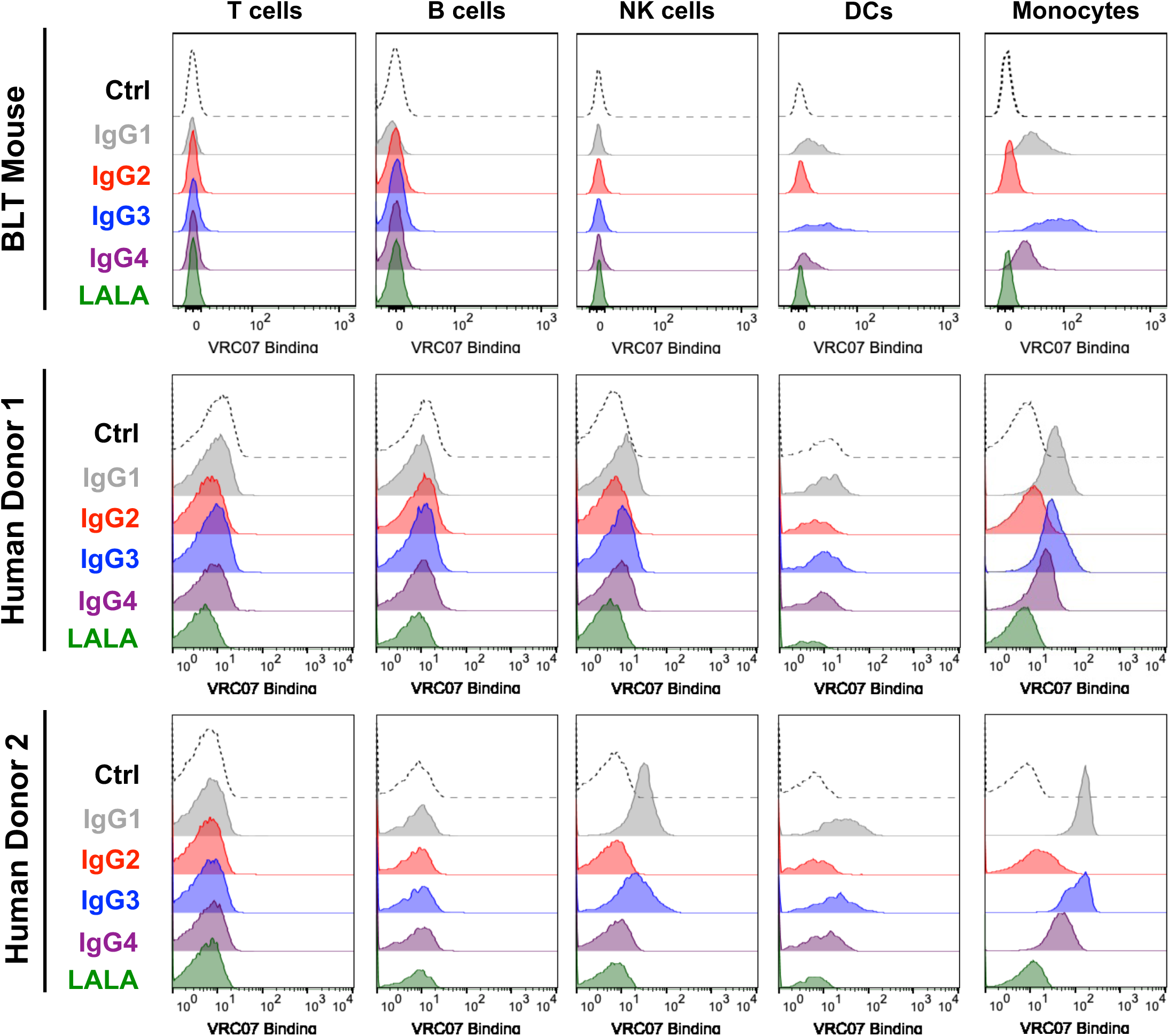
Binding of human immune cell subsets to VRC07 of each subclass, related to Figure 3. Human immune cells were isolated from the spleen of a representative BLT humanized mouse or the peripheral blood of two different human donors and mixed with AF647-labeled VRC07 Fc variants. Plots show mean fluorescence intensity measured by flow cytometry for T cells (CD45+, CD3+), B cells (CD45+, CD20+), NK cells (CD45+, CD3-, CD20-, CD14-, CD56+), DCs (CD45+, CD3-, CD20-, CD56-, CD16-, HLA-DR+, CD11c+), and monocytes (CD45+, CD3-, CD56-, CD14+).

**Figure S4.**
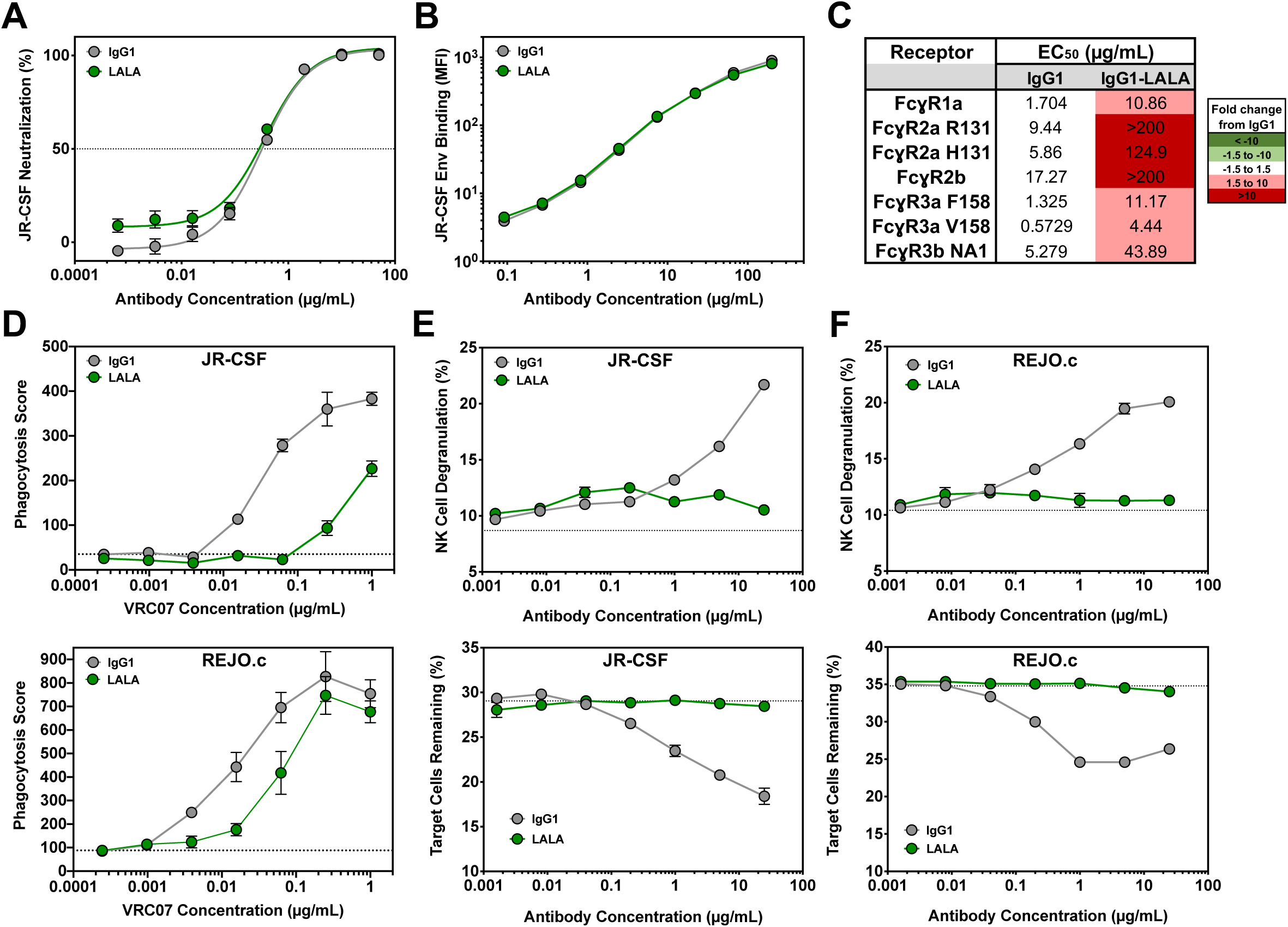
Impact of L234A, L235A (LALA) mutations on neutralization, Env binding and effector function activity of VRC07 IgG1, related to Figure 3. **(A)** *In vitro* neutralization activity of purified VRC07 IgG1 or LALA against HIV_JR-CSF_ as measured by TZM-bl neutralization assays. Error bars represent SEM. **(B)** Binding of VRC07 IgG1 or LALA to HIV_JR-CSF_ Env-expressing target cells as measured by flow cytometry with a pan-IgG-Fc detection reagent. Error bars represent SEM. **(C)** Half-maximal binding concentration (EC_50_) of cell surface-bound VRC07 IgG1 or LALA to purified fluorescent-labeled FcɣR proteins as measured by flow cytometry. The color of the box indicates the fold change in half-maximal binding compared to VRC07 IgG1. **(D)** THP-1 cell phagocytosis of HIV_JR-CSF_ (top) or HIV_REJO.c_ (bottom) Env-expressing target cells bound by VRC07 IgG1 or LALA as measured by flow cytometry. The dotted line indicates the average phagocytic score in the absence of antibody. Error bars represent SEM. **(E and F)** Ability of VRC07 IgG1 or LALA to mediate NK cell degranulation (top) as well as specific killing (bottom) of HIV_JR-CSF_ **(E)** or HIV_REJO.c_ **(F)** Env-expressing target cells as measured by flow cytometry. The dotted line indicates the average value in the absence of antibody. Error bars represent SEM.

**Figure S5.**
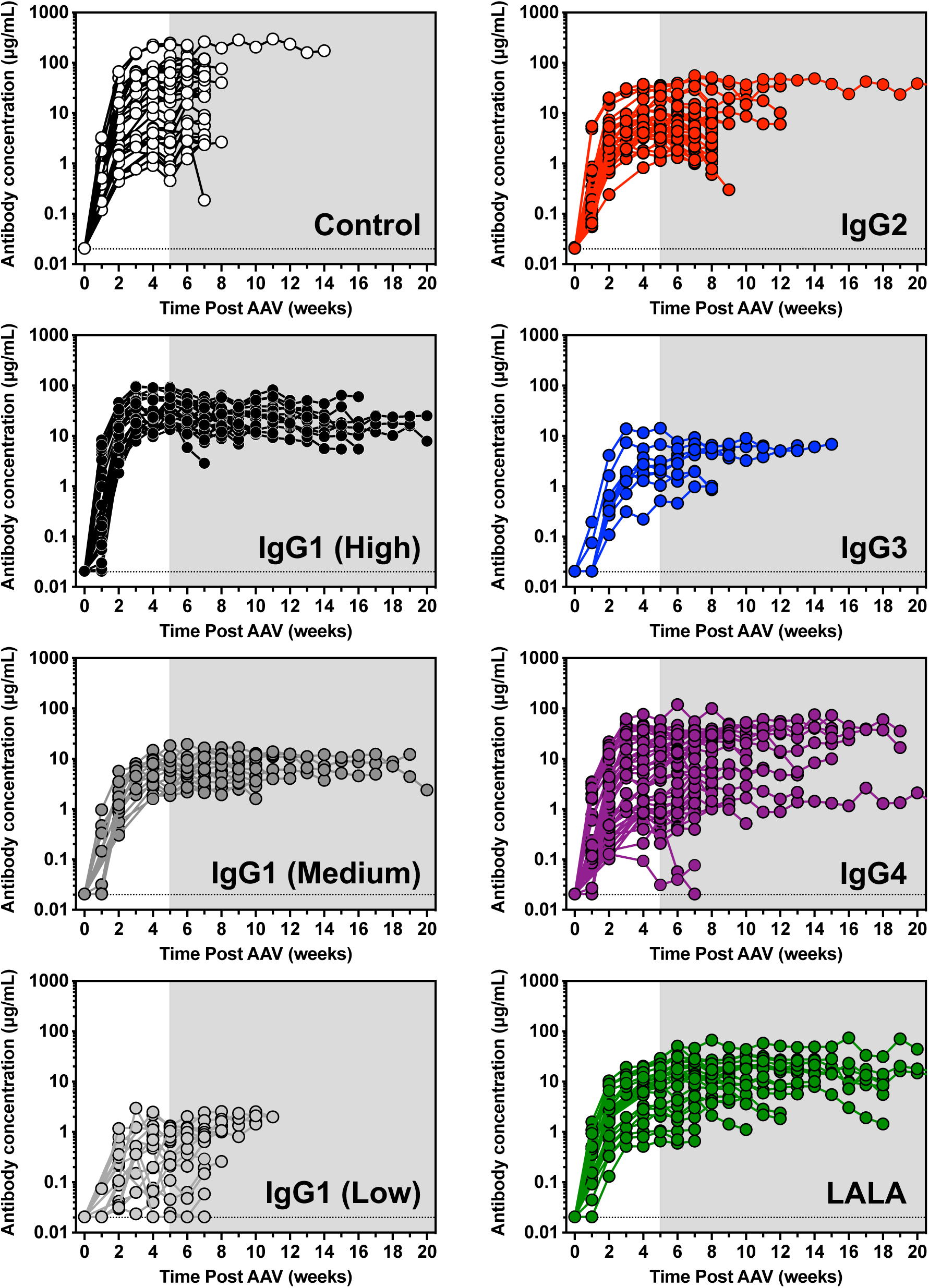
Antibody concentration of individual mice, related to Figure 3 and Figure 4. Antibody concentration (µg/mL) in the plasma of BLT humanized mice from the time of AAV injection until the time of HIV acquisition or death as measured by ELISA. The shaded region indicates the timing of weekly vaginal HIV challenges with 30 ng p24 HIV_REJO.c_, initiated at 5 weeks post AAV administration. The dotted line indicates the limit of detection (0.0206 µg/mL) for the ELISA.

**Figure S6.**
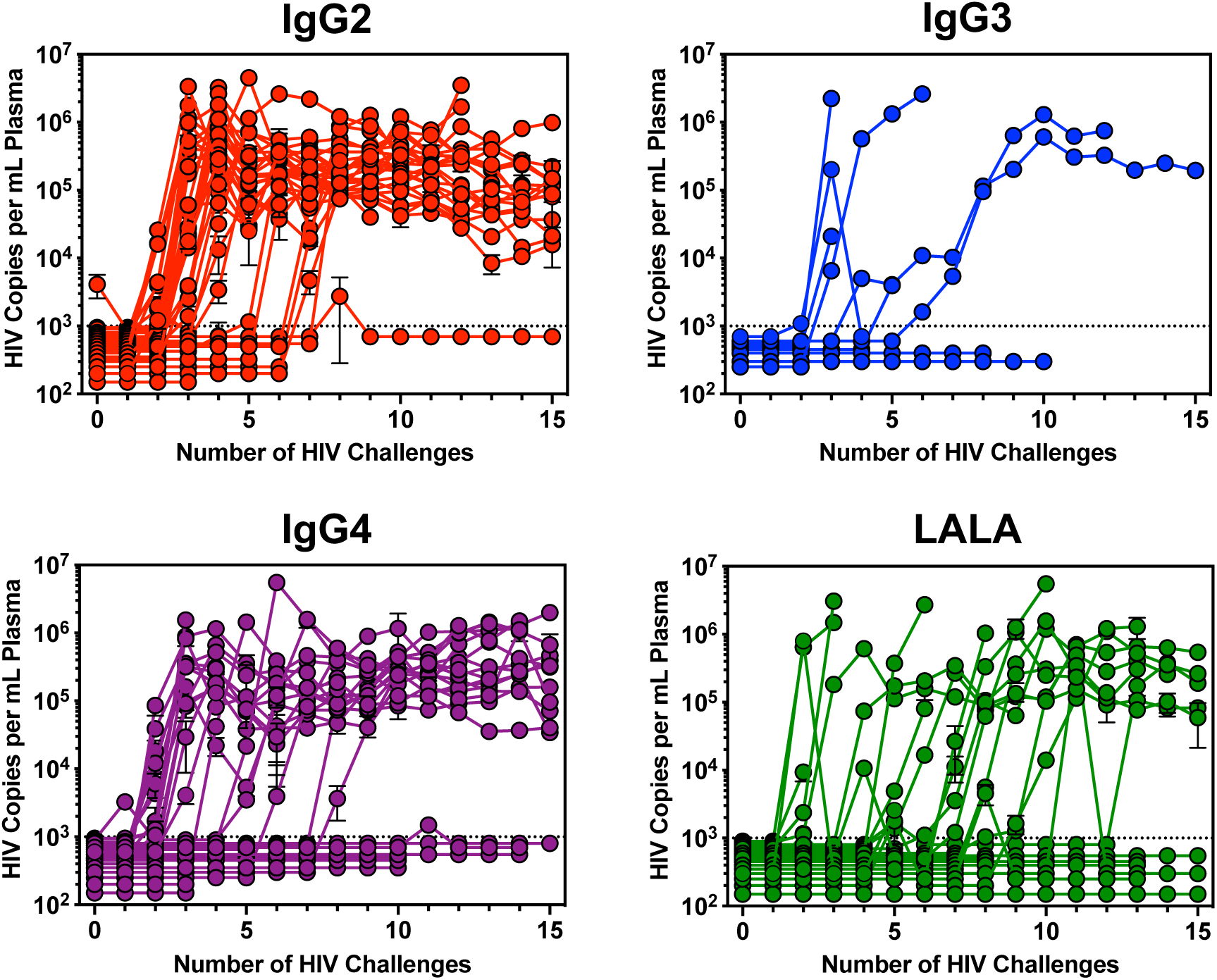
Viral Load of individual mice, related to Figure 3. Viral load in the plasma of BLT humanized mice during repetitive challenge with 30 ng p24 HIV_REJO.c_ as measured by qPCR. The dotted line indicates the limit of detection (1000 copies/mL) for the assay.

## METHODS

### RESOURCE AVAILABILITY

#### Lead Contact

Further information and requests for resources and reagents should be directed to and will be fulfilled by Alejandro Balazs.

#### Materials Availability

Plasmids generated in this study will be available through Addgene. Recombinant proteins and antibodies are available from their respective sources.

#### Data and Code Availability

This study did not generate sequence data or code. Data generated in the current study (including ELISA and neutralization) have not been deposited in a public repository but are available from the corresponding author upon request.

### EXPERIMENTAL MODEL AND SUBJECT DETAILS

#### Human subjects

The use of human tissues was approved by Partners Human Research Committee of MGH (Protocol 2012P000409). All donor tissues were obtained under an MGH Institutional Review Board (IRB) approved protocol for discarded, de-identified tissues from participants who agreed to use of donations for research. Participants received no compensation.

#### Mice

All experiments were done with approval from the Institutional Animal Care and Use Committee (IACUC) of the Massachusetts General Hospital and conducted in accordance with the regulations of the American Association for the Accreditation of Laboratory animal Care (AAALAC).

#### Cell lines

CEM.NKr cells were obtained from the NIH AIDS Reagent Program and engineered to express HIV_JR-CSF_ Env using lentiviral transduction. THP-1 cells were obtained from the American Type Culture Collection (ATCC).

### METHOD DETAILS

#### Construction and purification of VRC07 Fc variant antibodies

Class-switched VRC07 constructs were generated from human isotype backbone plasmids obtained from Addgene. The different human heavy chain constant regions were PCR amplified from the Addgene plasmids and inserted into the previously described AAV transfer vector ^11^ encoding the VRC07 heavy chain variable region and VRC07 kappa light chain. Sequencing was performed to confirm constant region insertion. Subsequently, each class-switched VRC07 gene was cloned into a third-generation self-inactivating lentiviral vector. Lentivirus was produced by co-transfection of 293T cells with the given VRC07 IgG lentiviral vector and two helper plasmids (pHDM-VSV-G and pHDM-Hgpm2). After 48 h, culture supernatant containing lentivirus was collected, filtered, and used to transduce fresh 293T cells in order to generate cell lines stably expressing VRC07 IgG antibodies. These cell lines were incubated in Freestyle 293 expression medium (Thermo Fisher Scientific) at 37°C for 10-15 days. VRC07 IgG antibodies were then purified from the culture supernatant using affinity chromatography with Pierce Protein A/G Agarose (Thermo Fisher Scientific) followed by size-exclusion chromatography using a Superdex 200 column (GE Healthcare) on an AKTA Purifier Fast Protein Liquid Chromatography system (GE Healthcare). Quantification of purified protein was performed using a NanoDrop 2000 Spectrophotometer (Thermo Fisher Scientific) and an IgG-specific ELISA with isotype-matched standards.

##### VRC07 antibody sequences

*VRC07 IgG1 heavy chain*: MATGSRTSLLLAFGLLCLPWLQEGSAQVRLSQSGGQMKKP GDSMRISCRASGYEFINCPINWIRLAPGKRPEWMGWMKPRGGAVSYARQLQGRVTMTRDMY SETAFLELRSLTSDDTAVYFCTRGKYCTARDYYNWDFEHWGQGTPVTVSSASTKGPSVFPLA PSSKSTSGGTAALGCLVKDYFPEPVTVSWNSGALTSGVHTFPAVLQSSGLYSLSSVVTVPSSS LGTQTYICNVNHKPSNTKVDKKVEPKSCDKTHTCPPCPAPELLGGPSVFLFPPKPKDTLMISRT PEVTCVVVDVSHEDPEVKFNWYVDGVEVHNAKTKPREEQYNSTYRVVSVLTVLHQDWLNGKE YKCKVSNKALPAPIEKTISKAKGQPREPQVYTLPPSRDELTKNQVSLTCLVKGFYPSDIAVEWE SNGQPENNYKTTPPVLDSDGSFFLYSKLTVDKSRWQQGNVFSCSVMHEALHNHYTQKSLSLS PG

##### VRC07 IgG2 heavy chain

MATGSRTSLLLAFGLLCLPWLQEGSAQVRLSQSGGQMKKP GDSMRISCRASGYEFINCPINWIRLAPGKRPEWMGWMKPRGGAVSYARQLQGRVTMTRDMY SETAFLELRSLTSDDTAVYFCTRGKYCTARDYYNWDFEHWGQGTPVTVSSASTKGPSVFPLA PCSRSTSESTAALGCLVKDYFPEPVTVSWNSGALTSGVHTFPAVLQSSGLYSLSSVVTVPSSN

FGTQTYTCNVDHKPSNTKVDKTVERKCCVECPPCPAPPVAGPSVFLFPPKPKDTLMISRTPEV TCVVVDVSHEDPEVQFNWYVDGVEVHNAKTKPREEQFNSTFRVVSVLTVVHQDWLNGKEYK CKVSNKGLPAPIEKTISKTKGQPREPQVYTLPPSREEMTKNQVSLTCLVKGFYPSDIAVEWESN GQPENNYNTTPPMLDSDGSFFLYSKLTVDKSRWQQGNVFSCSVMHEALHNHYTQKSLSLSPG

##### VRC07 IgG3 heavy chain

MATGSRTSLLLAFGLLCLPWLQEGSAQVRLSQSGGQMKKP GDSMRISCRASGYEFINCPINWIRLAPGKRPEWMGWMKPRGGAVSYARQLQGRVTMTRDMY SETAFLELRSLTSDDTAVYFCTRGKYCTARDYYNWDFEHWGQGTPVTVSSASTKGPSVFPLA PCSRSTSGGTAALGCLVKDYFPEPVTVSWNSGALTSGVHTFPAVLQSSGLYSLSSVVTVPSSS LGTQTYTCNVNHKPSNTKVDKRVELKTPLGDTTHTCPRCPEPKSCDTPPPCPRCPEPKSCDTP PPCPRCPEPKSCDTPPPCPRCPAPELLGGPSVFLFPPKPKDTLMISRTPEVTCVVVDVSHEDP EVQFKWYVDGVEVHNAKTKPREEQFNSTFRVVSVLTVLHQDWLNGKEYKCKVSNKALPAPIE KTISKTKGQPREPQVYTLPPSREEMTKNQVSLTCLVKGFYPSDIAVEWESSGQPENNYNTTPP MLDSDGSFFLYSKLTVDKSRWQQGNIFSCSVMHEALHNRFTQKSLSLSPG

##### VRC07 IgG4 heavy chain

MATGSRTSLLLAFGLLCLPWLQEGSAQVRLSQSGGQMKKP GDSMRISCRASGYEFINCPINWIRLAPGKRPEWMGWMKPRGGAVSYARQLQGRVTMTRDMY SETAFLELRSLTSDDTAVYFCTRGKYCTARDYYNWDFEHWGQGTPVTVSSASTKGPSVFPLA PCSRSTSESTAALGCLVKDYFPEPVTVSWNSGALTSGVHTFPAVLQSSGLYSLSSVVTVPSSS LGTKTYTCNVDHKPSNTKVDKRVESKYGPPCPSCPAPEFLGGPSVFLFPPKPKDTLMISRTPE VTCVVVDVSQEDPEVQFNWYVDGVEVHNAKTKPREEQFNSTYRVVSVLTVLHQDWLNGKEY KCKVSNKGLPSSIEKTISKAKGQPREPQVYTLPPSQEEMTKNQVSLTCLVKGFYPSDIAVEWES NGQPENNYKTTPPVLDSDGSFFLYSRLTVDKSRWQEGNVFSCSVMHEALHNHYTQKSLSLSL G

##### VRC07 kappa light chain

MATGSRTSLLLAFGLLCLPWLQEGSAEIVLTQSPGTLSLSPG ETAIISCRTSQYGSLAWYQQRPGQAPRLVIYSGSTRAAGIPDRFSGSRWGPDYNLTISNLESGD FGVYYCQQYEFFGQGTKVQVDIKRTVAAPSVFIFPPSDEQLKSGTASVVCLLNNFYPREAKVQ WKVDNALQSGNSQESVTEQDSKDSTYSLSSTLTLSKADYEKHKVYACEVTHQGLSSPVTKSF NRGEC

#### HIV production and quantification

Viruses were produced by transient transfection of 293T cells with 2 µg/mL of plasmid encoding HIV_NL4-3_, HIV_JR-CSF_, or HIV_REJO.c_ (NIH AIDS Reagent Program). After 48 hours, culture supernatants were collected, filtered with a 0.45 µm filter, and titered using both an HIV-1 p24 antigen capture assay (Leidos Biomedical Research) and 50% tissue culture infective dose (TCID_50_) assay on TZM-bl cells. TCID_50_ was calculated using the Spearman-Karber formula (Ramakrishnan, 2016).

#### In vitro neutralization assay

Neutralization assays were performed by combining 200 TCID_50_ of virus with three-fold serial dilutions of a given antibody. Virus-antibody mixtures were incubated at 37°C for 1 h. Following incubation, the mixtures were added to 96-well plates seeded with 10,000 TZM-bl cells per well 24 h prior to the assay. Media in each well was also supplemented with 75 µg/mL DEAE Dextran to facilitate viral infection. After 48 h at 37°C, cells were lysed using a previously-described luciferase buffer (Siebring-van Olst et al., 2013) and luciferase sexpression was measured on a Spectramax L microplate reader (Molecular Devices). Percent neutralization was calculated by subtracting the background luciferase signal of control wells (cells only) from the luciferase signal of experimental wells (cells with virus and antibody) and then dividing this value by the difference between virus control wells (cells with virus in the absence of antibody) and control wells with cells only.

#### Engineered CEM.NKr cells expressing cell-surface HIV Env

CEM.NKr cells were obtained from the NIH AIDS Reagent Program and engineered to express HIV_JR-CSF_ Env using lentiviral transduction. Briefly, the gene for membrane-bound HIV_JR-CSF_ gp150 was engineered for enhanced expression through human codon optimization, and replacement of the endogenous Env leader sequence with the leader sequence of human CD5 antigen (Haas et al., 1996). This engineered HIV_JR-CSF_ gp150 gene was synthesized (Integrated DNA Technologies) and cloned into a lentiviral vector (Mostoslavsky et al., 2006). The transgene was expressed under control of the human elongation factor 1 alpha promoter. An internal ribosome entry site (IRES) located downstream of the envelope transgene and drove cap-independent translation of the fluorescent protein, ZsGreen. Lentivirus was produced through co-transfection of 293T cells with the HIV_JR-CSF_ gp150 lentiviral vector and four helper plasmids (pHDM-VSV-G, pHDM-Hgpm2, pHDM-Tat1b, and pRC-CMV-Rev1b). After 48 h, cell culture supernatants were collected, filtered through a 0.45 µm filter, and used to transduce CEM.NKr cells. Following transduction, CEM.NKr cells were single-cell sorted using a FACSAria II SORP Flow Cytometer (BD Biosciences) and screened based on high Env and ZsGreen expression to generate a clonal target cell population.

#### *In vitro* effector function assays

Binding of VRC07 Fc variants to purified FcɣR proteins was measured using a flow-based assay as previously described (Brown et al., 2017). Either HIV_JR-CSF_ or HIV_REJO.c_ Env-expressing target cells were stained with three-fold serial dilutions of a given antibody for 30 min at room temperature followed by 0.65 µg/mL of a FcɣR detection reagent for 1 h at room temperature. The tetrameric detection reagents were generated immediately prior to use by combining biotinylated FcɣR proteins (Ackerman Laboratory) with a 1/4^th^ molar ratio of Streptavidin-Allophycocyanin (ProZyme) and mixing by inversion for 10 min. Free biotin was then added to a final concentration of 5 µM to completely block free streptavidin binding sites. The stained samples were analyzed on a Stratedigm S1300Exi Flow Cytometer. To determine the half-maximal effective concentration, or EC_50_, of each antibody, the mean fluorescence intensity (MFI) of APC and antibody concentrations were log transformed and fit with a non-linear regression using variable slope function in GraphPad Prism.

To measure ADCP, a bead-based assay was used as previously described (Ackerman et al., 2011; Brown et al., 2017). Briefly, purified HIV_JR-CSF_ gp120 protein was biotinylated using EZ-Link Sulfo-NHS-LC-Biotin (Thermo Fisher Scientific) and excess biotin was removed using a Zeba spin desalting column (Thermo Fisher Scientific). Biotinylated antigen was then conjugated to fluorescent neutravidin-labeled microspheres. The gp120-conjugated beads were incubated with five-fold serial dilutions of a given antibody for 2 h at 37°C in 96-well plates. Following incubation, 25,000 THP-1 cells were added to each well in a final volume of 200 µL and plates were incubated at 37°C overnight. Cells were then analyzed on a Stratedigm S1300Exi flow cytometer and a phagocytosis score was calculated by multiplying the percent of bead-positive THP-1 cells by the mean fluorescence intensity of the bead-positive cell population.

In addition to the bead-based ADCP assay, a cell-based assay was performed using ZsGreen+ Env-expressing CEM.NKr cell lines. Briefly, these target cells were mixed with parental CEM.NKr cells at a 1:1 ratio followed by three-fold serial dilutions of a given antibody. After 15 min at room temperature, THP-1 cells stained with CellTrace Violet were added at a 1:1 effector:target ratio and the co-cultures were incubated at 37°C overnight. The following day, cells were washed with 1X PBS and then fixed with 4% PFA for 20 min. Cells were analyzed on a Stratedigm S1300Exi flow cytometer and phagocytosis was measured as the percent of ZsGreen+ THP-1 cells.

To measure ADCC, HIV_JR-CSF_ or HIV_REJO.c_ Env-expressing CEM.NKr cells were combined with parental CEM.NKr cells at a 1:1 ratio. The mixed target population was then incubated with five-fold serial dilutions of a given antibody for 30 min at room temperature. Following incubation, NK cells isolated from buffy coats using an EasySep human NK cell enrichment kit (StemCell Technologies) were added to the mixed target population at a 1:1 effector:target ratio. Prior to addition, NK cells were stained with CellTrace Violet for 20 min at room temperature. PE/Cy7-conjugated anti-human CD107a antibody (BioLegend) was added to each effector-target cell mixture at a final concentration of 2-3 µg/mL and plates were incubated at 37°C for 6 h. Following incubation, cells were stained with LIVE/DEAD Fixable Far Red dye (Thermo Fisher Scientific), fixed with BD Cytofix fixation buffer (BD Biosciences), and analyzed on a Stratedigm S1300Exi flow cytometer. Degranulation of NK cells was measured as the percent of live, CD107a+, CellTrace Violet+ cells. Specific cell killing was measured by reduction in the percentage of the live, ZsGreen+ CEM.NKr target cells.

#### Generation of humanized mice for *in vivo* challenge studies

Immunodeficient NOD/SCID IL2Rgamma^null^ (NSG) mice were obtained from the Jackson Laboratory. To generate human PBMC-engrafted NSG mice, frozen human PBMCs (AllCells) were thawed and expanded in RPMI-1640 medium (Sigma-Aldrich) supplemented with 10% fetal bovine serum (VWR International), 1% L-glutamine (Thermo Fisher Scientific), 10 mM HEPES (Corning Inc), 1X non-essential amino acids (Corning Inc), 1X sodium pyruvate (Corning Inc), 50µM beta-mercaptoethanol (Agilent Technologies), 1% penicillin-streptomycin (Corning Inc), and stimulated for T-cell expansion with 5 µg/mL phytohemagglutinin (Sigma-Aldrich) and 10 ng/mL human IL-2. After 10 days of *in vitro* expansion, 4 million cells were administered to NSG mice by intraperitoneal injection in a 300 µL volume. Mice were rested two weeks following cell administration to allow for engraftment.

Bone marrow-liver-thymus (BLT) humanized mice were generated by the Human Immune System Mouse Program at the Ragon Institute of MGH, MIT and Harvard. Briefly, 6-8 week old female NSG mice were transplanted with human liver and thymus tissue under the kidney capsule and injected intravenously with 100,000 CD34+ cells isolated from liver tissue by AutoMACS. Mice were rested ten weeks following surgery to allow for recovery and engraftment.

#### Flow cytometry of mouse samples for FcɣR expression profiling

Ten weeks after surgery, blood samples were taken from BLT mice by retro-orbital bleeding and centrifuged at 1,150 x g for 5 min at room temperature to separate plasma from the cell pellets. Plasma was removed and frozen at -80°C for subsequent analysis. The cell pellets were resuspended in 1.1 mL of 1x RBC lysis buffer (BioLegend) and incubated on ice for 10 min. After red blood cell lysis, each sample was pelleted at 1,150 x g in a centrifuge for 5 min at room temperature and then stained with 50 µL of an antibody cocktail containing 1:50 diluted anti-human CD45-AF700 (Biolegend, clone HI30), 1:100 diluted anti-human CD3-FITC (BioLegend, clone UCHT1), 1:25 diluted anti-human CD19-PE/Cy7 (BioLegend, clone SJ25C1), 1:25 diluted anti-human CD56-BV570 (BioLegend, clone HCD56), 1:25 diluted anti-human CD14-PE (BioLegend, clone HCD14), 1:25 diluted anti-human CD11c-BV421 (BioLegend, clone Bu15), 1:50 diluted anti-human HLA-DR-BV510 (BioLegend, clone L243), 1:50 diluted anti-human CD16-APC/Cy7 (BioLegend, clone B73.1), 1:50 diluted anti-human CD32-APC (BioLegend, clone FUN-2), and 1:50 diluted anti-human CD64-PE/DAZZLE594 (BioLegend, clone 10.1) in PBS supplemented with 2% FBS (PBS+) for 30 min on ice. Samples were then analyzed on a Stratedigm S1300Exi flow cytometer. T cells were defined as a CD45+, CD3+ population. B cells were defined as a CD45+, CD19+ population. NK cells were defined as a CD45+, CD3-, CD19-, HLA-DR-, CD56+ population. Myeloid cells were defined as a CD45+, CD3, CD19-, CD11c+, HLA-DR+ population. The frequency FcɣR expression (i.e. CD64, CD32, and CD16) was then assessed for each cell population.

#### Flow cytometry of mouse samples for VRC07 subclass staining of myeloid populations

Spleens were harvested from BLT mice at 25 weeks or 34 weeks post-surgery and washed twice with complete RPMI at 500 x g. HuPBMCs were isolated from whole blood by Histopaque density gradient centrifugation (Sigma-Aldrich). Cells were stained with 1.75µg/ml of VRC07-AlexaFluor647 conjugated isotypes, VRC07-IgG1, VRC07-IgG2, VRC07-IgG3, VRC07-IgG4 and VRC07-IgG1-LALA at 4°C for 30 minutes in PBS. Cells were washed with PBS supplemented with 2% FBS (PBS+) and then stained with 50µl of an antibody cocktail containing 1:50 diluted anti-human CD45-AF700 (BioLegend, clone HI30), 1:50 diluted anti-human CD3-BV605 (BioLegend, clone UCHT1), 1:50 diluted anti-human CD20-BV650 (BioLegend, clone 2H7), 1:50 diluted anti-human HLA-DR-BV570 (BioLegend, clone L243), 1:50 diluted anti-human CD15-PE-Cy5 (BioLegend, clone W6D3), 1:50 diluted anti-human CD13-PerCP-Cy5.5 (BioLegend, clone WM15), 1:50 diluted anti-human CD16-BV785 (BioLegend, clone 3G8), 1:50 diluted anti-human CD14-PE/Cy7 (BioLegend, clone M5E2), 1:50 diluted anti-human CD11b-BV421 (BioLegend, clone LM2), 1:50 diluted anti-human CD11c-FITC (BD, clone B-ly6), 1:50 diluted anti-human CD56-PE (BioLegend, clone QA17A16), and 1:100 diluted LIVE/DEAD-Near-IR in PBS+ for 30 minutes at 4°C. Samples were then washed and fixed in 4%PFA and analyzed on a Stratedigm S1300Exi flow cytometer. T cells were defined as a CD45+, CD3+ population. B cells were defined as a CD45+, CD20+ population. NK cells were defined as a CD45+, CD3-, CD14-, CD11c-, CD56+ population. Classical monocytes were defined as CD45+, CD3-, CD56-, CD15-, CD16-, CD14+ population. Macrophages were defined as a CD45+, CD3-, CD56-, CD14+, CD16+, CD64+, CD11b+ population. Neutrophils were defined as a CD45+, CD3-, CD56-, CD15+, CD13+ population. DCs were defined as a CD45+, CD3-, CD56-, CD14-, CD16-, HLA-DR+, CD11c+ population. VRC07 engagement with specific cell types was shown as histograms and the mean fluorescence intensity (MFI) of AF647 was calculated for each isotype staining.

#### AAV virus production, quantification, and administration

AAV2/8 encoding a given antibody was produced via transient transfection of 293T cells and purification from culture supernatant via PEG precipitation and cesium chloride ultracentrifugation as previously described (Balazs et al., 2011). AAV titers were then determined by qPCR. Briefly, AAV aliquots were serially diluted and incubated with 1.2 U of Turbo DNase (Invitrogen) for 30 min at 37°C. 5 µL of DNase-treated virus was used in a 15 µl qPCR reaction with PerfeCTa SYBR Green SuperMix, ROX (Quanta Biosciences) and primers designed against the CMV enhancer (5’-AACGCCAATAGGGACTTTCC and 3’-GGGCGTACTTGGCATATGAT). Samples were run in duplicate on a QuantStudio 12K Flex Real-Time PCR system (Applied Biosystems) with the following cycling conditions: 50°C for 2 min, 95°C for 10 min, followed by 50 cycles of 95°C for 15 s and 60°C for 60 s. Virus titer was determined by comparison with a standard curve generated using AAV transfer vector plasmid DNA. Intramuscular injections of titered AAV were performed as previously described (Balazs et al., 2011). Briefly, aliquots of virus were thawed on ice and diluted to the desired dose in a 40 µL volume. A single injection of 40 µL was administered into the gastrocnemius muscle of NSG mice or BLT humanized mice using a 28 G insulin syringe.

#### HIV challenge of humanized mice

HuPBMC-NSG mice were challenged with 280 TCID_50_ HIV_NL4-3_ in a 50 µL volume two weeks after humanization with PBMCs. IV challenge was administered through retro-orbital injection of the venous sinus using a 28 G insulin syringe. BLT mice were challenged with 30 ng p24 of HIV_REJO.c_ in a 20 µL volume. Vaginal challenge was performed non-abrasively by placing isoflurane-anesthetized mice in a supine position and elevating the posterior of the animal before shallow insertion of the pipette loaded with virus into the vaginal vault. After viral administration, mice were maintained in a supine position for 5 min to prevent loss of the virus. This vaginal challenge protocol was repeated weekly until the conclusion of the experiment. For all mouse experiments, mice were bled weekly to determine CD4+ T cell counts, plasma antibody concentrations, and plasma viral loads.

#### Flow cytometry of blood samples

Blood samples were taken from mice by retro-orbital bleeding and centrifuged at 1,150 x g for 5 min at room temperature to separate plasma from the cell pellets. Plasma was removed and frozen at -80°C for subsequent analysis. The cell pellets were resuspended in 1.1 mL of 1x RBC lysis buffer (BioLegend) and incubated on ice for 10 min. After red blood cell lysis, each sample was pelleted at 1,150 x g in a centrifuge for 5 min at room temperature and then stained with 50 µL of an antibody cocktail containing 1:100 diluted anti-human CD3-FITC (BioLegend, clone UCHT1), 1:100-diluted anti-human CD4-PE (BioLegend, clone RPA-T4), and 1:100 diluted anti-human CD8-APC (BioLegend, clone RPA-T8) in PBS supplemented with 2% FBS (PBS+) for 30 min on ice. Samples were then analyzed on a Stratedigm S1000 flow cytometer. Samples were first gated by CD3 expression before determining the ratio of CD4 to CD8 within this subset. Samples containing fewer than 20 CD3+ events were excluded from the analysis.

#### Antibody quantification by ELISA

For detection of gp120-binding IgG in mouse sera, ELISA plates were coated with 0.2 µg per well of purified HIV_JR-CSF_ gp120 protein for 1 h at room temperature. Plates were then blocked with 1% BSA (KPL) in Tris-buffered saline (TBS) for at least 30 min at room temperature or overnight at 4°C. Mouse serum samples were diluted in TBS plus Tween 20 (TBST) containing 1% BSA (KPL) and incubated on the plate for 1 h at room temperature, followed by a 30 min incubation with HRP-conjugated goat anti-human IgG Fc antibody at a 1:2500 dilution (Bethyl, A80-104A). Detection was completed using the TMB Microwell Peroxidase Substrate System (KPL) and a VersaMax microplate reader (Molecular Devices). Samples were compared to a standard curve generated using purified VRC07 antibodies of the same IgG subclass.

#### Viral load quantification by quantitative RT-PCR

Viral RNA was extracted from mouse sera using the QIAGEN QIAamp Viral RNA Mini Kit. Each RNA sample was treated with 2 U of Turbo DNase (Invitrogen) for 30 min at 37°C, followed by 15 min at 75°C for heat inactivation. To quantify viral RNA, 10 µL of the DNase-treated sample was used in a 20 µL RT-qPCR reaction with qScript XLT one-step RT-qPCR ToughMix low-ROX (Quanta Biosciences), a TaqMan probe (5’-/56-FAM/CCCACCAAC/ZEN/AGGCGGCCTTAACTG/3IABkFQ/-3’) (IDT) and primers designed to target the Pol gene of HIV_REJO.c_ (5’-CAATGGCCCCAATTTCATCA and 3’-GAATGCCGAATTCCTGCTTGA) or HIV_NL4-3_ (5’ CAATGGCAGCAATTTCACCA and 3’ GAATGCCAAATTCCTGCTTGA). Samples were run in triplicate on a QuantStudio 12K Flex Real-Time PCR system (Applied Biosystems) with the following cycle conditions: 50°C for 10 min, 95°C for 3 min followed by 55 cycles of 95°C for 3 s and 60°C for 30 s. Virus titer was determined by comparison with a standard curve generated using RNA extracted from a serially-diluted mixture of commercially-titered viral stock and pure mouse serum. For all viral strains, the limit of detection was 1,000 HIV genome copies per mL.

### QUANTIFICATION AND STATISTICAL ANALYSIS

An infection event was defined as two consecutive positive viral loads, and the week of the first positive sample was defined as the time of HIV acquisition. The survival data was fit to a cox proportional hazards regression model using the coxph function from the survival package v3.2-7 (https://CRAN.Rproject.org/package=survival) in R v4.0.2 (R Core Team 2020). All mice that survived at least 7 weeks after AAV administration were included in the analysis.

## REFERENCES

1. Ackerman, M.E., Moldt, B., Wyatt, R.T., Dugast, A.-S., McAndrew, E., Tsoukas, S., Jost, S., Berger, C.T., Sciaranghella, G., Liu, Q., et al. (2011). A robust, high-throughput assay to determine the phagocytic activity of clinical antibody samples. Journal of Immunological Methods 366, 8–19.

2. Ackerman, M.E., Das, J., Pittala, S., Broge, T., Linde, C., Suscovich, T.J., Brown, E.P., Bradley, T., Natarajan, H., Lin, S., et al. (2018). Route of immunization defines multiple mechanisms of vaccine-mediated protection against SIV. Nat Med 24, 1590–1598.

3. Alter, G., Yu, W.-H., Chandrashekar, A., Borducchi, E.N., Ghneim, K., Sharma, A., Nedellec, R., McKenney, K.R., Linde, C., Broge, T., et al. (2020). Passive Transfer of Vaccine-Elicited Antibodies Protects against SIV in Rhesus Macaques. Cell 183, 185–196.e14.

4. Balazs, A.B., Chen, J., Hong, C.M., Rao, D.S., Yang, L., and Baltimore, D. (2011). Antibody-based protection against HIV infection by vectored immunoprophylaxis. Nature 481, 81–84.

5. Balazs, A.B., Ouyang, Y., Hong, C.M., Chen, J., Nguyen, S.M., Rao, D.S., An, D.S., and Baltimore, D. (2014). Vectored immunoprophylaxis protects humanized mice from mucosal HIV transmission. Nature Medicine 20, 296–300.

6. Banerjee, K., Klasse, P.J., Sanders, R.W., Pereyra, F., Michael, E., Lu, M., Walker, B.D., and Moore, J.P. (2010). IgG Subclass Profiles in Infected HIV Type 1 Controllers and Chronic Progressors and in Uninfected Recipients of Env Vaccines. AIDS Research and Human Retroviruses 26, 445–458.

7. Bournazos, S., Klein, F., Pietzsch, J., Seaman, M.S., Nussenzweig, M.C., and Ravetch, J.V. (2014). Broadly Neutralizing Anti-HIV-1 Antibodies Require Fc Effector Functions for In Vivo Activity. Cell 158, 1243–1253.

8. Brown, E.P., Dowell, K.G., Boesch, A.W., Normandin, E., Mahan, A.E., Chu, T., Barouch, D.H., Bailey-Kellogg, C., Alter, G., and Ackerman, M.E. (2017). Multiplexed Fc array for evaluation of antigen-specific antibody effector profiles. Journal of Immunological Methods 443, 33–44.

9. Casazza, J.P., Cale, E.M., Narpala, S., Novik, L., Yamshchikov, G., Lin, B.C., Pandey, J.P., McDermott, A.B., Roederer, M., Balazs, A.B., et al. (2021). DURABLE HIV-SPECIFIC bNAb PRODUCTION IN HUMANS AFTER AAV8 MEDIATED GENE TRANSFER. Conference on Retroviruses and Opportunistic Infections.

10. Chu, T.H., Crowley, A.R., Backes, I., Chang, C., Tay, M., Broge, T., Tuyishime, M., Ferrari, G., Seaman, M.S., Richardson, S.I., et al. (2020). Hinge length contributes to the phagocytic activity of HIV-specific IgG1 and IgG3 antibodies. PLoS Pathog 16, e1008083.

11. Corey, L., Gilbert, P.B., Juraska, M., Montefiori, D.C., Morris, L., Karuna, S.T., Edupuganti, S., Mgodi, N.M., deCamp, A.C., Rudnicki, E., et al. (2021). Two Randomized Trials of Neutralizing Antibodies to Prevent HIV-1 Acquisition. N Engl J Med 384, 1003–1014.

12. Crowley, A.R., and Ackerman, M.E. (2019). Mind the Gap: How Interspecies Variability in IgG and Its Receptors May Complicate Comparisons of Human and Non-human Primate Effector Function. Front. Immunol. 10, 697.

13. Deal, C., Balazs, A.B., Espinosa, D.A., Zavala, F., Baltimore, D., and Ketner, G. (2014). Vectored antibody gene delivery protects against Plasmodium falciparum sporozoite challenge in mice. Proceedings of the National Academy of Sciences of the United States of America 111, 12528–12532.

14. Denton, P.W., Estes, J.D., Sun, Z., Othieno, F.A., Wei, B.L., Wege, A.K., Powell, D.A., Payne, D., Haase, A.T., and Garcia, J.V. (2008). Antiretroviral pre-exposure prophylaxis prevents vaginal transmission of HIV-1 in humanized BLT mice. PLoS Medicine 5, e16.

15. Haas, J., Park, E.C., and Seed, B. (1996). Codon usage limitation in the expression of HIV-1 envelope glycoprotein. Current Biology : CB 6, 315–324.

16. Hangartner, L., Beauparlant, D., Rakasz, E., Nedellec, R., Hozé, N., McKenney, K., Martins, M.A., Seabright, G.E., Allen, J.D., Weiler, A.M., et al. (2021). Effector function does not contribute to protection from virus challenge by a highly potent HIV broadly neutralizing antibody in nonhuman primates. Sci. Transl. Med. 13, eabe3349.

17. Hessell, A.J., Hangartner, L., Hunter, M., Havenith, C.E.G., Beurskens, F.J., Bakker, J.M., Lanigan, C.M.S., Landucci, G., Forthal, D.N., Parren, P.W.H.I., et al. (2007). Fc receptor but not complement binding is important in antibody protection against HIV. Nature 449, 101–104.

18. Hezareh, M., Hessell, A.J., Jensen, R.C., van de Winkel, J.G., and Parren, P.W. (2001). Effector function activities of a panel of mutants of a broadly neutralizing antibody against human immunodeficiency virus type 1. Journal of Virology 75, 12161–12168.

19. Hughes, J.P., Baeten, J.M., Lingappa, J.R., Magaret, A.S., Wald, A., de Bruyn, G., Kiarie, J., Inambao, M., Kilembe, W., Farquhar, C., et al. (2012). Determinants of Per-Coital-Act HIV-1 Infectivity Among African HIV-1–Serodiscordant Couples. The Journal of Infectious Diseases 205, 358–365.

20. Janeway, C.A., Travers, P., Walport, M., Shlomchik, M.J., Jr, C.A.J., Travers, P., Walport, M., and Shlomchik, M.J. (2001). Immunobiology (Garland Science).

21. Julg, B., Tartaglia, L.J., Keele, B.F., Wagh, K., Pegu, A., Sok, D., Abbink, P., Schmidt, S.D., Wang, K., Chen, X., et al. (2017). Broadly neutralizing antibodies targeting the HIV-1 envelope V2 apex confer protection against a clade C SHIV challenge. Science Translational Medicine 9, eaal1321.

22. Moldt, B., Rakasz, E.G., Schultz, N., Chan-Hui, P.-Y., Swiderek, K., Weisgrau, K.L., Piaskowski, S.M., Bergman, Z., Watkins, D.I., Poignard, P., et al. (2012a). Highly potent HIV-specific antibody neutralization in vitro translates into effective protection against mucosal SHIV challenge in vivo. Proceedings of the National Academy of Sciences of the United States of America 109, 18921–18925.

23. Moldt, B., Shibata-Koyama, M., Rakasz, E.G., Schultz, N., Kanda, Y., Dunlop, D.C., Finstad, S.L., Jin, C., Landucci, G., Alpert, M.D., et al. (2012b). A nonfucosylated variant of the anti-HIV-1 monoclonal antibody b12 has enhanced FcγRIIIa-mediated antiviral activity in vitro but does not improve protection against mucosal SHIV challenge in macaques. Journal of Virology 86, 6189–6196.

24. Mostoslavsky, G., Fabian, A.J., Rooney, S., Alt, F.W., and Mulligan, R.C. (2006). Complete correction of murine Artemis immunodeficiency by lentiviral vector-mediated gene transfer. Proceedings of the National Academy of Sciences of the United States of America 103, 16406– 16411.

25. Neidich, S.D., Fong, Y., Li, S.S., Geraghty, D.E., Williamson, B.D., Young, W.C., Goodman, D., Seaton, K.E., Shen, X., Sawant, S., et al. (2019). Antibody Fc effector functions and IgG3 associate with decreased HIV-1 risk. Journal of Clinical Investigation 129, 4838–4849.

26. Om, K., Paquin-Proulx, D., Montero, M., Peachman, K., Shen, X., Wieczorek, L., Beck, Z., Weiner, J.A., Kim, D., Li, Y., et al. (2020). Adjuvanted HIV-1 vaccine promotes antibody-dependent phagocytic responses and protects against heterologous SHIV challenge. PLoS Pathog 16, e1008764.

27. Parsons, M.S., Lee, W.S., Kristensen, A.B., Amarasena, T., Khoury, G., Wheatley, A.K., Reynaldi, A., Wines, B.D., Hogarth, P.M., Davenport, M.P., et al. (2018). Fc-dependent functions are redundant to efficacy of anti-HIV antibody PGT121 in macaques. The Journal of Clinical Investigation.

28. Pegu, A., Borate, B., Huang, Y., Pauthner, M.G., Hessell, A.J., Julg, B., Doria-Rose, N.A., Schmidt, S.D., Carpp, L.N., Cully, M.D., et al. (2019). A Meta-analysis of Passive Immunization Studies Shows that Serum-Neutralizing Antibody Titer Associates with Protection against SHIV Challenge. Cell Host & Microbe 26, 336–346.e3.

29. Ramakrishnan, M.A. (2016). Determination of 50% endpoint titer using a simple formula. World Journal of Virology 5, 85–86.

30. Richardson, S.I., Lambson, B.E., Crowley, A.R., Bashirova, A., Scheepers, C., Garrett, N., Abdool Karim, S., Mkhize, N.N., Carrington, M., Ackerman, M.E., et al. (2019). IgG3 enhances neutralization potency and Fc effector function of an HIV V2-specific broadly neutralizing antibody. PLoS Pathog 15, e1008064.

31. Shangguan, S., Ehrenberg, P.K., Geretz, A., Yum, L., Kundu, G., May, K., Fourati, S., Nganou-Makamdop, K., Williams, L.D., Sawant, S., et al. (2021). Monocyte-derived transcriptome signature indicates antibody-dependent cellular phagocytosis as a potential mechanism of vaccine-induced protection against HIV-1. ELife 10, e69577.

32. Shattock, R.J., and Moore, J.P. (2003). Inhibiting sexual transmission of HIV-1 infection. Nat Rev Microbiol 1, 25–34.

33. Siebring-van Olst, E., Vermeulen, C., de Menezes, R.X., Howell, M., Smit, E.F., and van Beusechem, V.W. (2013). Affordable luciferase reporter assay for cell-based high-throughput screening. Journal of Biomolecular Screening 18, 453–461.

34. Sips, M., Krykbaeva, M., Diefenbach, T.J., Ghebremichael, M., Bowman, B.A., Dugast, A.-S., Boesch, A.W., Streeck, H., Kwon, D.S., Ackerman, M.E., et al. (2016). Fc receptor-mediated phagocytosis in tissues as a potent mechanism for preventive and therapeutic HIV vaccine strategies. Mucosal Immunol 9, 1584–1595.

35. Sok, D., and Burton, D.R. (2018). Recent progress in broadly neutralizing antibodies to HIV. Nat Immunol 19, 1179–1188.

36. Stoddart, C.A., Maidji, E., Galkina, S.A., Kosikova, G., Rivera, J.M., Moreno, M.E., Sloan, B., Joshi, P., and Long, B.R. (2011). Superior human leukocyte reconstitution and susceptibility to vaginal HIV transmission in humanized NOD-scid IL-2Rγ(-/-) (NSG) BLT mice. Virology.

37. Sun, Z., Denton, P.W., Estes, J.D., Othieno, F.A., Wei, B.L., Wege, A.K., Melkus, M.W., Padgett-Thomas, A., Zupancic, M., Haase, A.T., et al. (2007). Intrarectal transmission, systemic infection, and CD4+ T cell depletion in humanized mice infected with HIV-1. The Journal of Experimental Medicine 204, 705–714.

38. Vidarsson, G., Dekkers, G., and Rispens, T. (2014). IgG Subclasses and Allotypes: From Structure to Effector Functions. Front. Immunol. 5.

39. Visciano, M.L., Tagliamonte, M., Tornesello, M.L., Buonaguro, F.M., and Buonaguro, L. (2012). Effects of adjuvants on IgG subclasses elicited by virus-like Particles. J Transl Med 10, 4.

40. Yi, J.S., Rosa-Bray, M., Staats, J., Zakroysky, P., Chan, C., Russo, M.A., Dumbauld, C., White, S., Gierman, T., Weinhold, K.J., et al. (2019). Establishment of normative ranges of the healthy human immune system with comprehensive polychromatic flow cytometry profiling. PLoS ONE 14, e0225512.

